# Structural Insights into the Assembly and Regulation of 2’-*O* RNA Methylation by SARS-CoV-2 nsp16/nsp10

**DOI:** 10.1101/2024.12.19.628950

**Authors:** Anurag Misra, R. Rahisuddin, Manish Parihar, Shailee Arya, Thiruselvam Viswanathan, Nathaniel Jackson, Shan Qi, Siu-Hong Chan, Reuben S. Harris, Luis Martinez-Sobrido, Yogesh K. Gupta

**Affiliations:** Greehey Children’s Cancer Research Institute, University of Texas Health Science Center, San Antonio, TX 78229, USA; Department of Biochemistry and Structural Biology, University of Texas Health Science Center, San Antonio, TX 78229, USA; Texas Biomedical Research Institute, San Antonio, TX, 78227, USA; New England Biolabs, 240 County Road, Ipswich, MA, 01938, USA; Howard Hughes Medical Institute, University of Texas Health Science Center, San Antonio, TX 78229, USA

## Abstract

2’-*O*-ribose methylation of the first transcribed base (adenine or A_1_ in SARS-CoV-2) of viral RNA mimics the host RNAs and subverts the innate immune response. How nsp16, with its obligate partner nsp10, assembles on the 5’-end of SARS-CoV-2 mRNA to methylate the A_1_ has not been fully understood. We present a ∼ 2.4 Å crystal structure of the heterotetrameric complex formed by the cooperative assembly of two nsp16/nsp10 heterodimers with one 10-mer Cap-1 RNA (product) bound to each. An aromatic zipper-like motif in nsp16 and the N-terminal regions of nsp10 and nsp16 orchestrate an oligomeric assembly for efficient methylation. The front catalytic pocket of nsp16 stabilizes the upstream portion of the RNA while the downstream RNA remains unresolved, likely due to its flexibility. An inverted nsp16 dimer extends the positively charged surface area for longer RNA to influence the catalysis. Additionally, a non-specific nucleotide-binding pocket on the backside of nsp16 plays a critical role in catalysis, further contributing to its enzymatic activity.

## Introduction

Coronaviruses (CoVs) enzymatically modify 2’-OH of ribose sugar of the first transcribed nucleotide (N_1_) to protect their mRNA from degradation and escape from the host innate immune recognition^1^. The 5’-end of N_1_ base (usually A_1_ in CoVs, A_1_/G_1_ in other RNA viruses) is enzymatically modified with the sequential attachment of an RNA cap and its subsequent methylation by S-adenosyl-*L*-methionine (SAM)-dependent methyltransferases^2, 3^. The capping of SARS-CoV-2 mRNA includes a combinatorial action of the nidovirus RdRp-associated nucleotidyltransferase (NiRAN) domain of nonstructural protein 12 (nsp12); nsp13, a terminal GTP hydrolase; nsp9, a substrate for AMPylation activity during RNA cap formation^4^ (**Fig. 1A**). Consistent with its predecessors, the 2’-OH methylation of A_1_ base in SARS-CoV-2 is embodied in nsp16, which acts as a SAM-dependent 2’-*O*-methyltransferase (2’-*O*-MTase) in conjunction with its noncatalytic stimulator nsp10^5–13^. The nsp16/nsp10 complex converts the status of the mRNA cap from Cap-0 (^m7^GpppA) to Cap-1 (^m7^GpppAm). Previous studies on SARS-CoV, MERS-CoV, and even SARS-CoV-2 proteins suggest nsp16/nsp10 heterodimer to be an active 2’-*O*-MTase^7–13^. The Cap-1 formation by nsp16/nsp10 can avert the recognition of SARS-CoV-2 mRNA by melanoma differentiation-associated protein 5 (MDA5) and shields it from type I interferon (IFN-I)-induced antiviral response^14^. Consistently, genetic perturbation of nsp16 activity leads to the induction of IFN-I via the MDA5 RNA sensor^14–17^.

**Fig. 1.**
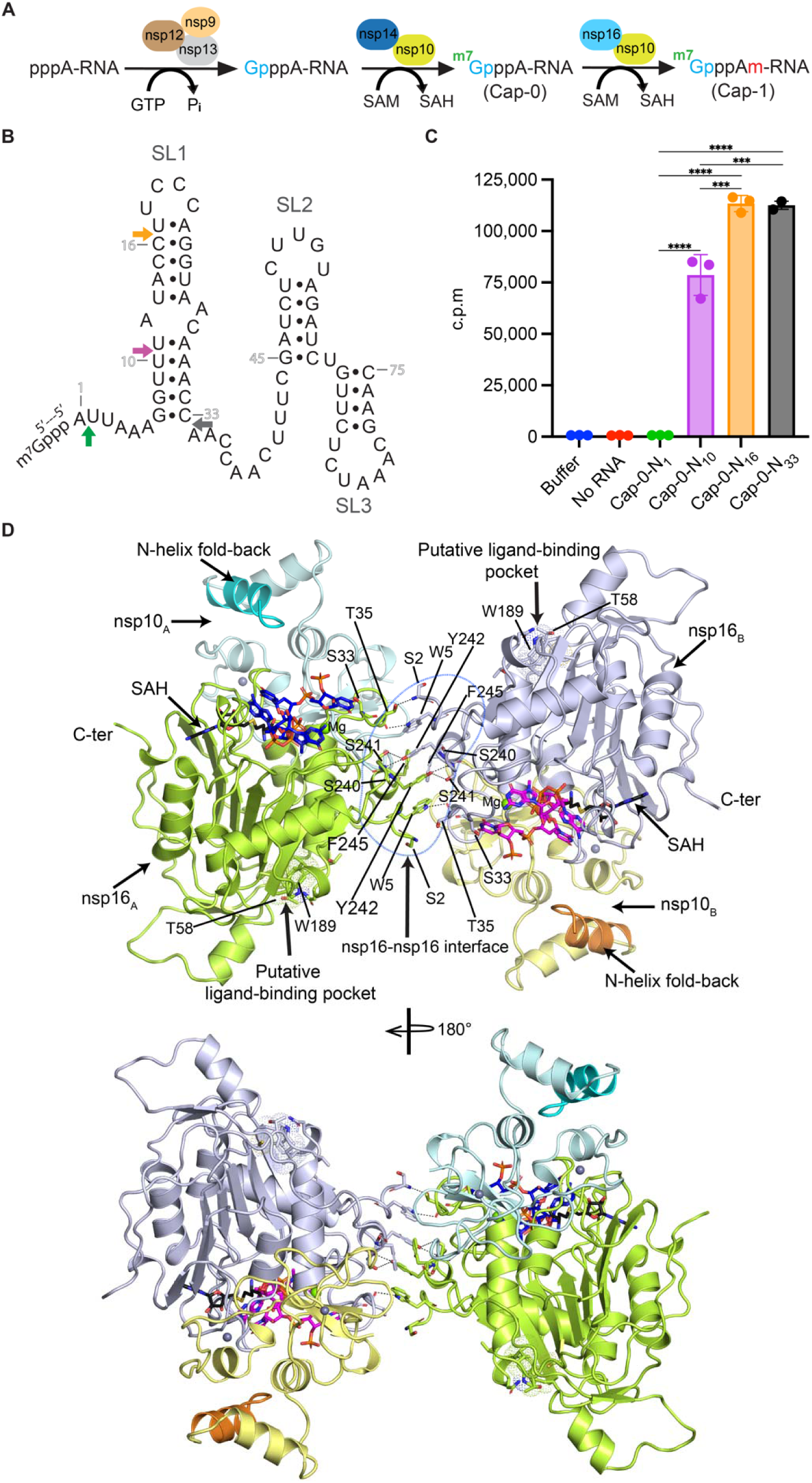
Target RNA length and overall structure. **A.** Schematic of RNA capping and modification. **B.** Sequence and secondary structure of the 5’ leader sequence of SARS-CoV-2 RNA depicting the three stem-loop (SL1, SL2, and SL3) regions. Cap-0-RNA 10-mer, 16-mer, and 33-mer were used to assess the substrate preference of nsp16/nsp10 for methyltransferase activity. Statistical analysis was performed using one-way ANOVA with multiple comparisons. ‘Buffer’ and ‘No RNA’ groups were excluded from the test. **** p < 0.0001, *** p < 0.001. **C.** A radiometric assay exhibiting enzyme activity on RNA substrates with varied length (n=3). **D.** Final structure of nsp16/nsp10 heterotetramer in the presence of a 10-mer Capped RNA (Cap-1); first three nucleotides (blue and magenta sticks) and SAH (black sticks) were refined with full occupancy. Two inverted nsp16/nsp10 heterodimers are stabilized by an aromatic zipper-like interface.

Clinical studies have reported the lack or a significant delay in the production of IFN-I and IFN-III in patients with severe COVID-19 disease^18^. Hence, the ablation of nsp16 activity should trigger an immune response to CoV infection to limit pathogenesis^1, 15^. Genomic disruption of SARS-CoV nsp16 reduces the synthesis of viral RNA replication^16, 19^. Absence of 2’-*O*-methylation activity strongly enhances IFN-I responses against viral infections^1, 14, 15^ and attenuates viral replication^16, 20^. Thus, nsp16/nsp10 complex has emerged as an attractive therapeutic target^17, 21^. Consistently, short peptides of nsp10 encompassing the region that interacts with nsp16 could block the nsp16/nsp10 interface, thereby reducing viral replication and pathogenesis *in vitro* and animal models^22^.

According to a conventional model of 2’-*O*-methylation in CoVs, nsp16 interacts with nsp10 through a conserved hydrophobic interface and stabilizes the nsp16/nsp10 heterodimer, potentially extending the RNA binding groove. The allosteric stimulation of the 2’-*O*-MTase activity of nsp16 by nsp10, as a universally conserved mechanism for all CoVs, including SARS-CoV-2, was postulated to occur through the canonical nsp16/nsp10 heterodimeric interface^8, 9, 12,13^. Our previous work postulated the existence of a putative nucleotide-binding site on the opposite face of nsp16^8^. However, the mechanics of nsp16/nsp10 assembly on the 5’-end of mRNAs longer than six nucleotides (nts) and the precise role of the alternative ligand binding site are not fully understood, mainly due to the inability of co-crystallizing the complex with a longer Cap-0 RNA. Also, apart from extending the RNA binding groove of nsp16, nsp10’s role was unclear, specifically how nsp10 may engage with the RNA. In addition to viral RNAs, nsp16, independent of nsp10, can engage with the spliceosome machinery by interacting with the splice site of U1 and the branch point site of U2 small nuclear RNAs to globally suppress the host mRNA splicing during viral infection^23^. Thus, a better understanding of the RNA binding and structural assembly of nsp16 and the stimulator of its enzymatic activity nsp10 is required.

Here, we report a ∼ 2.4 Å structure of a heterotetrameric complex of SARS-CoV-2 nsp16/nsp10 in the presence of a 10-mer Cap-1 RNA with four out of ten nucleotides resolved and S-adenosyl homocysteine (SAH), a byproduct of the methylation reaction. This structure reveals an aromatic zipper motif in nsp16, which promotes the formation of new interfaces between two opposing nsp16/nsp10 heterodimers and together with a ∼ 132° flip of an N-terminal helix of nsp10, coordinates the assembly of the nsp16/nsp10 heterotetramer, an inverted dimer of nsp16/nsp10 heterodimer on RNA, for efficient 2’-*O*-methylation, a mark essential for host immune restriction. In addition, we now validated the putative ligand-binding pocket at the opposite face (back-side of the catalytic surface) of nsp16, which we and others previously identified as a non-specific nucleotide-binding site^8–10^. This pocket has the propensity to accommodate a variety of nucleotides and small ligands, including adenosine, m^7^GDP, m^7^GTP, m^7^GpppA, and MES^8–10, 12^. Our mutagenesis data indicate that this pocket modulates the 2’-*O*-methylation activity of nsp16/nsp10, though the precise mechanism by which this distant pocket impacts enzymatic activity remains unclear due to the absence of electron density for the downstream RNA, which may extend to this pocket.

## Results

### CoV RNA substrate preference

CoVs have the longest genome among all positive-strand RNA viruses. The first 394 nucleotides at the 5’-end of CoVs RNA fold into an array of secondary structural elements (stem loop; SL1-7)^24^. This region harbors a leader sequence of ∼ 75 nts at the 5’-end, which plays an essential role in the discontinuous synthesis of nine sub-genomic RNAs in the β-CoVs, including SARS-CoV-2^25^. The leader sequence folds into three distinct stem-loops (SL1-3) with 6 nts, including target adenine (A_1_), remain unpaired at the 5’-terminus (**Fig. 1B**). The 3’-end of the leader sequence overlaps with SL3 and contains a 6 nts long (CUAAAC) transcription regulatory sequence (TRS-L) that fuses with the genome just upstream of the coding sequence for each transcription unit [TRS-B (body)] during transcription^25^. On the other hand, the SL1 plays an equally important role in the viral life cycle^26^. Previous structural studies for 2′-*O*-MTase activity in SARS-CoV-2 were performed with Cap-0 analog (^m7^GpppA) or a short 6-mer Cap-0 RNA (^m7^GpppAUUAAA)^8, 9, 12, 13^.

To determine the length of RNA required by SARS-CoV-2 nsp16/nsp10 for optimal 2’-*O*-MTase activity, we designed and synthesized three cognate Cap-0 RNA oligonucleotides encompassing varied lengths of SARS-CoV-2 mRNA: the 10-mer (^m7^GpppAUUAAAGGUU) includes additional four nucleotides downstream of the six unpaired bases; the 16-mer (^m7^GpppAUUAAAGGUUUAUACC) includes ten unpaired bases; and the 33-mer (^m7^GpppAUUAAAGGUUUAUACCUUCCCAGGUAACAAACC) has the entire SL1 sequence that follows the 6 unpaired bases (**Fig. 1B**). We used an established radiometric assay to measure methyl transfer from _1_H^3^-labeled SAM (S-adenosyl-*L*-methionine) to the 2’-OH of the first adenine base (N_1_) of RNA in the presence of Mg^2+^ as a cofactor in the reaction buffer. We detected negligible methylation with Cap-0 analog (Cap-0-N_1_) but robust methylation with a 10-mer Cap-0 RNA (Cap-0-N_10_) as a substrate, corroborating the previous observation (Cap-0-N_6_)^13^. Adding 6 more nucleotides to the 3’-end (Cap-0-N_16_) further increased nsp16/nps10 activity. However, the longest 33-mer RNA encompassing the entire SL1 region (Cap-0-N_33_) did not enhance methylation beyond what was achieved with the 16-mer RNA (**Fig. 1C**). The preference for longer substrates with partial or complete SL1 sequence highlights the importance of nucleotides from N_1_ to N_16_ for optimal methylation of Cap-0 RNA of SARS-CoV-2. A similar observation was made for a critical enzymatic step during RNA capping that precedes 2’-*O*-methylation (i.e. RNAylation of nsp9 by NiRAN domain of nsp12) where a 10-mer 5’-pppRNA corresponding to the first 10 bases of the leader sequence yielded robust RNAylation activity^4^.

### Crystallization and structure determination of SARS-CoV-2 nsp16/nsp10 complex

Nsp16/nsp10 heterodimeric structures in complex with short 6-mer Cap-0 RNA (^m7^GpppAUUAAA) have been reported^13^. However, these structures were resolved by soaking Cap-0 analogs or short Cap-0 RNAs into apo nsp16/nsp10 crystals. Since nsp16/nsp10 shows higher methylation in the presence of longer (>10 nts Cap-0 RNA), we attempted to co-crystallize the SARS CoV-2 nsp16/nsp10 complex in the presence of longer Cap-0 RNA oligos. After rigorous attempts, we obtained co-crystals of SARS-CoV-2 nsp16/nsp10 in the presence of a 10-mer Cap-0 RNA (^m7^GpppA-N_10_; ^m7^GpppA_1_•U_2_•U_3_•A_4_•A_5_•A_6_•G_7_•G_8_•U_9_•U_10_) and S-adenosyl homocysteine (SAH), a byproduct of the methylation reaction. Our best crystals diffract X-rays to ∼ 2.4 Å resolution with synchrotron radiation (NECAT 24ID-C beamline at Advanced Photon Source, Chicago). Previously, the nsp16/nsp10 heterodimer in complex with soaked SAH/SAM and Cap-0 analogs or short Cap-0 RNAs (^m7^GpppA-N_6_) was crystallized in a trigonal space group (P3_1_21 or P3_2_21)^8, 9, 12, 13^. The new crystals of nsp16/nsp10 grown in the presence of the ^m7^GpppA-N_10_ RNA belong to a different space group (monoclinic, P2_1_) with unit cell dimensions of a = 111.75 Å, b = 58.89 Å, c = 117.02 Å, and α = γ = 90°, β=95.48° (**Table 1**). Based on Vm calculations, two nsp16/nsp10/RNA complexes (related by a 2-fold non-crystallographic symmetry) are present in the crystallographic asymmetric unit. We solved the structure by molecular replacement using the SARS-CoV-2/SAM/^m7^GpppA complex (PDB ID: 6WKS)^8^ structure as a template (**Table 1**).

**Table 1.**
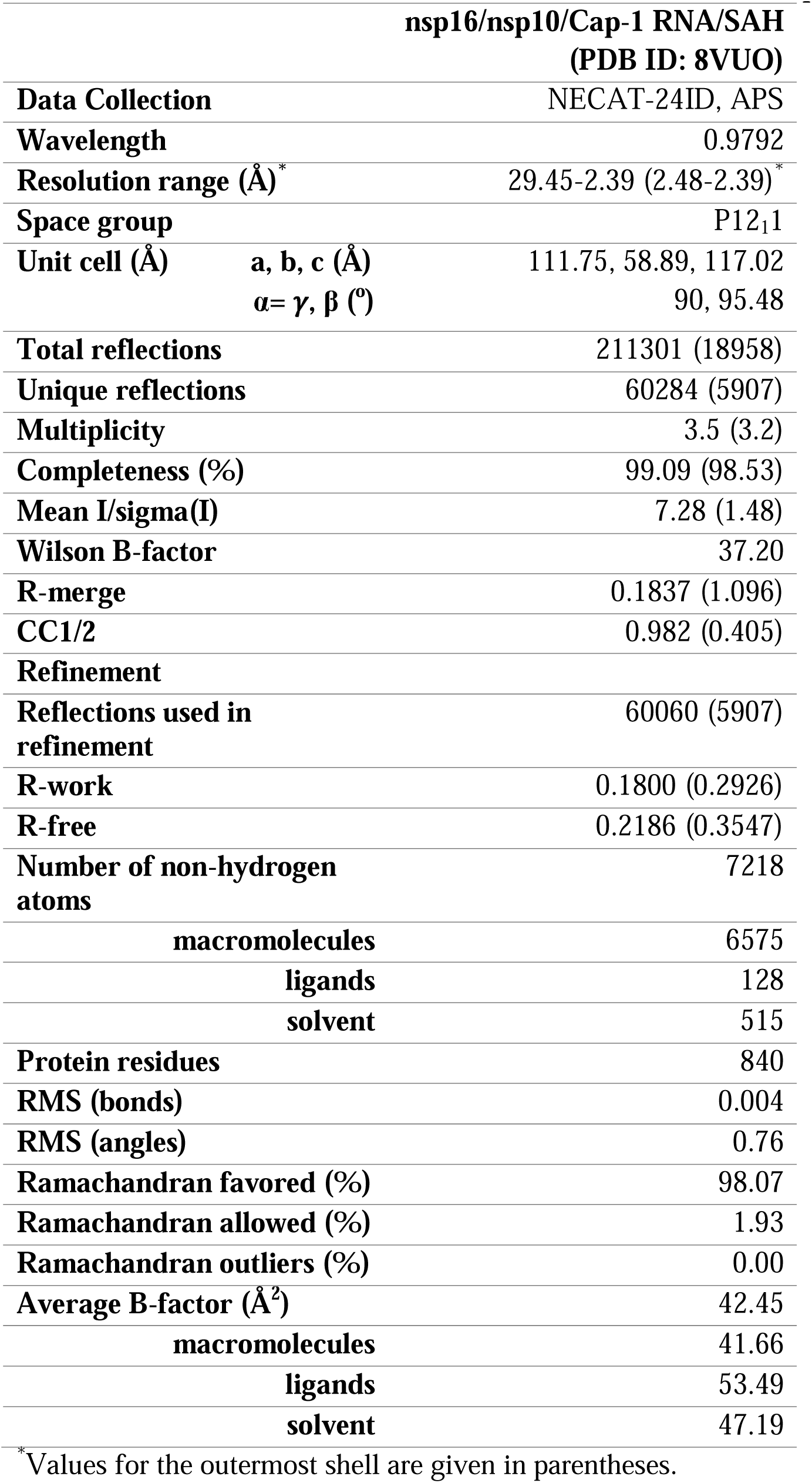
Data collection and refinement statistics (molecular replacement).

We observed clear density for two nsp16 and two nsp10 molecules. The final refined model contains two chains of nsp16 (chain A and chain C, each chain contains aa S2 - V294) and two chains of nsp10 (chain B and chain D, each containing aa V7 - Q132). All chains encompass well-resolved regions, except for the terminal residues: S1 at the N-terminus and L295 - N298 at the C-terminus of nsp16, as well as A1 - E6 at the N-terminus and L133 - Q139 at the C-terminus of nsp10 (**Fig. 1D**). Among the two chains of 10-mer Cap-0 RNA (chain E and F), we resolved the electron density for the first three nucleotides (^m7^Gppp**<u>A_1_</u>**•U_2_•U_3_•) and the PO_4_ group of the fourth nucleotide, A_4_ **(Fig. S1A)**. The remaining nucleotides (A_4_•A_5_•A_6_•G_7_•G_8_•U_9_•U_10_) showed residual/untraceable electron density, hence not built, suggesting high flexibility of the 3’-end of the RNA. Though we used substrate RNA (Cap-0) for co-crystallization, however, in the electron density, we clearly resolved RNA in its product form (Cap-1, ^m7^Gppp**<u>A_m1_</u>**•U_2_•U_3_•), suggesting an *in crystallo* transfer of a methyl group from the co-purified SAM bound to nsp16/nsp10. The electron density for the 2’-*O* methyl and SAH moieties was unambiguously identified in the difference (Fo-Fc) electron density maps (**Fig. S1B**). Co-purification of SAM or SAH with recombinant SAM-dependent viral RNA methyltransferases from the expression host *E. coli* is a commonly observed phenomenon^9, 12, 27^.

### Overall structure of nsp16/nsp10/ ^m7^GpppA_m1_ RNA complex

A structural comparison of nsp16/nsp10 in heterotetrameric complex with previously reported structures of SARS-CoV-2 nsp16/nsp10 with Cap-0 analog (PDB ID: 6WKS)^8^, with Cap-0 short RNA (PDB ID: 7JYY), and Cap-1 short RNA (PDB ID: 7L6R^13^, revealed high structural similarity with the root mean square deviations of 0.331 Å (371 Cα), 0.346 Å (371 Cα), and 0.304 Å (363 Cα), respectively (**Fig. S1C, D**). Thus, the core structures of individual nsp16/nsp10 heterodimers, including their gate loops and catalytic pocket regions critical for substrate binding and catalysis, remain structurally conserved. However, the heterotetrameric structure reveals several astonishing features, which may resolve a paradox in the 2’-*O*-methylation field. The nsp10 binding to nsp16 is well known to stimulate RNA binding and catalytic activity of nsp16^5, 7, 8, 13^ (**Fig. 2A, B**). This canonical nsp16-nsp10 interface shares a buried surface area of ∼ 931 Å^2^ at the protein-protein interface^8^. Interestingly, the heterotetrameric assembly of two nsp16, two nsp10, and two RNA molecules create a new stable nsp16-nsp16 protein-protein interface between the two opposing nsp16/nsp10 heterodimers with an additional ∼ 492 Å^2^ buried surface area (**Fig. 2C - E**). In this regard, the new nsp16-nsp16 interface we revealed in the heterotetramer appears to be as extensive and stable as the nsp16-nsp10 interface, indicating its functional significance.

**Fig. 2.**
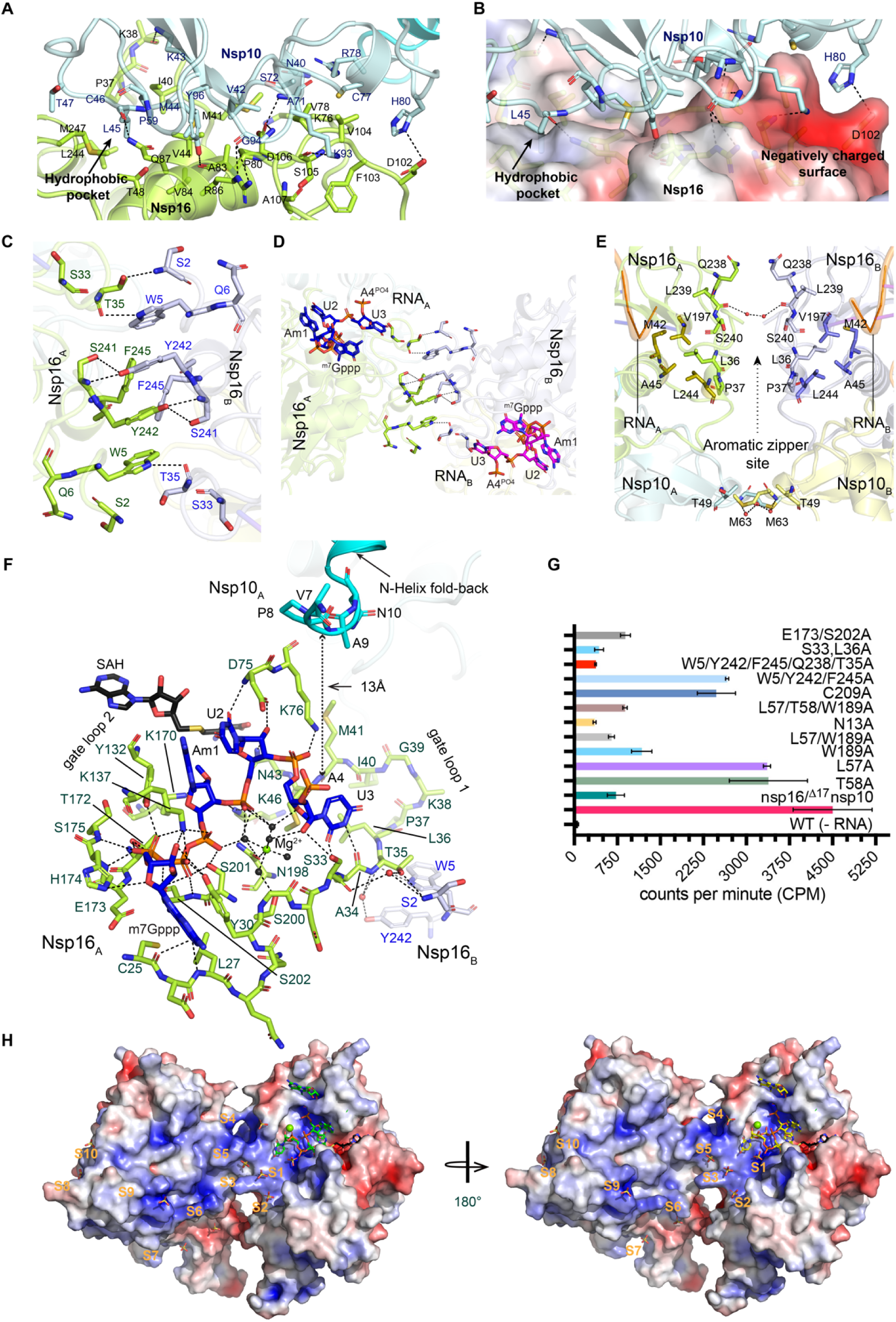
Nsp16-nsp10 (heterodimeric) and nsp16-nsp16 (heterotetrameric) interfaces. **A.** canonical nsp16-nsp10 interface in stick mode. **B.** electrostatic surface of nsp16 at the canonical nsp16-nsp10 interface. **C.** nsp16-nsp16 interface in the heterotetramer. Aromatic zipper-like motif. **D.** Aromatic zippers holding the tetramer and RNA together in *trans*. **E.** Residues at nsp16-nsp16 interface other than aromatic zipper. **F.** Protein-RNA interface showing interaction network. **G.** A radiometric assay shows reduced methyltransferase activity to varying degree of nsp16/nsp10 mutants (n=3). **H.** Electrostatic surface of the tetrameric nsp16/nsp10 complex displays a continuous positive surface formed by two opposing nsp16 protomers; probably, helpful to facilitate the interactions with an elongated RNA chain. SO4 ions bound to this surface in previously solved structures (PDB ID: 6WQ3, 6WRZ and 6WVN) are labeled as S1-S10.

Of note, Tyr242 and Phe245 residues of one nsp16 protomer form π-π stacking interactions with their counterparts from the second nsp16 protomer. At the same time, Tyr242 extends this stacking interaction to Trp5 within the same protomer (**Fig. 2C**). While Ser2 of nsp16_B_ forms h-bond with Thr35 of nsp16_A_ whereas another such pair at the interface remains at the van der Waals distance. The backbone carbonyl of Thr35 of nsp16_A_ forms h-bonding with the aromatic ring of Trp5 from nsp16_B,_ and the same bonding is present for another such pair at the interface (**Fig. 2C, D**). As such, the upstream portion of RNA (around the catalytic center) is held tightly by respective nsp16 protomers and strengthened by newly developed interface residues (Ser33, Ala34 and Thr35) (**Fig. 2D-F**), while the middle portion of RNA (starting from A_4_, poor electron density, hence not built) appears to be sandwiched between the N-helix of nsp10 of the same heterodimer and the N-terminus of nsp16 from the opposing heterodimer (**Fig. 2F**). Such a unique cooperative assembly of the catalytic subunit (nsp16) and their obligate noncatalytic partner (nsp10), at least to our knowledge, has not been observed for any 2’-*O*-MTase.

Another striking feature is the orientation of the N-terminal region (aa 1 - 17) of nsp10 in each heterodimer. This region could not be built in previous structures due to a lack of electron density in the absence of RNA. In the new structure, part of this region termed as N-helix (aa 11-19) (cyan cartoon in **Fig. 1D, 2F**) was well resolved in electron density maps. The N-terminus of nsp10 runs toward the RNA A_4_ base (only the phosphate of A_4_ was built), indicating its role in stabilizing the RNA. We also observed a Mg^2+^ ion that stabilizes the protein-RNA interface near the catalytic center of each nsp16 with octahedral geometry (**Fig. 2F, S1E**). The Mg^2+^-nsp16 interaction network includes Ser33, a residue mutated to Arg in nsp16 of a SARS-CoV-2 strain circulated during the pandemic in New York City^28^. We evaluated the impact of this residue on 2’-*O*-methylation. In the tetrameric complex, Ser33 is located within the gate loop1 region that interacts with the RNA cap and bases downstream to target A_1_ (**Fig. S1E, S2**). As such, the terminal region of RNA cap (m7Gppp), the target A_1_ base, and the next two bases (U_2_ and U_3_) of RNA are surrounded by two flexible gate loops 1 (aa 20 - 40) and 2 (aa 132 - 143) (**Fig. 2F, S2**). Specifically, Ser33 stabilizes the RNA by direct h-bonds with the 2’-OH of the U_3_ base (**Fig. 2F, Fig. S1E**), which breaks the stacking with the U_2_ base by rotating ∼ 180° into the pocket formed by gate loop 1 residues that include Ser33 and Leu36. Interestingly, this critical region (Ser33 – Leu36) of gate loop 1 of nsp16_A_ is sandwiched between RNA from one side and the N-terminal region of the other nsp16_B_ at nsp16-nsp16 dimer interface to facilitate the assembly of the two nsp16 protomers. Replacement of these two residues with Ala significantly reduced methylation activity (**Fig. 2G)**.

### Canonical nsp16/nsp10 interface

The canonical nsp16/nsp10 interface is stabilized by conserved hydrophobic and electrostatic interactions (**Fig. 2A, B, Fig. S2**). Although this interface was revealed in Cap-0 analog (PDB ID: 6WKS) and Cap-0 short RNA bound (PDB ID: 7JYY) nsp16/nsp10 structures, the buried surface area (BSA) was increased in the tetrameric complex likely due to the presence of longer RNA, e.g., ∼ 938 Å^2^ (Leu45^nsp10^, ∼ 169.57 Å^2^; His80^nsp10^, ∼ 66.87 Å^2^) in the current structure compared to ∼ 910 Å^2^ (Leu45^nsp10^, ∼ 157.60 Å^2^; His80^nsp10^, ∼ 42.11 Å^2^) in the Cap-0 6-mer RNA structure (PDB ID:7JYY), and ∼ 906 Å^2^ (Leu45^nsp10^, ∼ 160.44 Å^2^; His80^nsp10^, ∼ 46.18 Å^2^) in the Cap-0 analog-bound structure (PDB ID: 6WKS). Thus, longer RNA strengthens the canonical nsp16-nsp10 heterodimeric interface with a major contribution by the increase in the BSA for invariant Leu45 and His80 (**Fig. 2A, B)**.

As such, hydrophobic interactions dominate the canonical nsp16-nsp10 interface, especially at one end around the critical residue Leu45 of nsp10. The other end, however, is stabilized by charged electrostatic interactions around His80 from nsp10 (**Fig. 2A, B**). Leu45 acts as a wedge and buries in the hydrophobic pocket constituted by Pro37, Val44, Thr48, Leu244, and Met247, all from nsp16. The increase in the buried surface area at the canonical interface in the current complex occurs due to an inward movement of His80^nsp10^ toward Asp102^nsp16^, thereby promoting the compactness of this interface. Consistently, the Cα-Cα distance observed for the His80-Asp102 pair is ∼ 9.4Å (current structure) compared to ∼ 10.4Å (Cap-0 6-mer, PDB ID: 7JYY)^13^, and ∼ 10.0Å (Cap-0 analog, PDB ID: 6WKS)^8^ (**Fig. 2A, B**).

### A new heterotetrameric interface in the nsp16/nsp10 complex

The tetrameric assembly of the two nsp16-nsp10 dimers reveals a new nsp16-nsp16 homo-dimeric interface that includes an aromatic zipper-like motif (Trp5, Tyr242, and Phe245) further supported by Ser2, Ser33, Ala34, and Thr35. Trp5, Tyr242, and Phe245 form the aromatic zipper from the two opposing nsp16 (**Fig. 2C**). The interface residues belong to two 3_10_ helices, η1 (Ser2, and Trp5) and η5 (Ser241, Tyr242, and Phe245) at the N- and C-terminal ends of nsp16, respectively (**Fig. S2**). In addition, an important interface residue of gate loop 1 of nsp16_A_ is Thr35, which forms a hydrogen bond with the indole ring of Trp5 from the opposing nsp16_B_. An ∼ 1.3 Å shift in Thr35 (Cα) compared to the previous nsp16/nsp10 structures (PDB ID: 6WKS: ∼ 0.7 Å, 7JYY: ∼ 1.3 Å, 7L6R: ∼ 0.8 Å) facilitates such interactions (**Fig. S1D**). Also, the indole ring of Trp5 undergoes a 180° flip, and such an arrangement facilitates H-bonding with Thr35 and the new nsp16-nsp16 interface (**Fig. 2C, D, S1D**). Aromatic zippers are well known to promote protein oligomerization through aromatic stacking and hydrogen bonding^29, 30^.

Additional hydrophobic, van der Waals, and electrostatic interactions at this nsp16-nsp16 interface mediated by Gln238, Leu239, Ser240, Leu36, and Pro37 further support the aromatic zipper-like interface, thereby extending the total buried surface area by ∼ 66.89 Å^2^ (**Fig. 2E**). As such, side chains of Leu36 and Pro37 locate towards the hydrophobic core constituted by residues from α2 (Met42, Ala45), η4 (Val197), and η5 (Leu244) in both nsp16 (**Fig. 2E**). Ser33, Ala34, Thr35, Leu36 and Pro37 from gate loop 1 of nsp16_A_ stabilize the U_3_ base of RNA from one side and the new aromatic zipper interface residues of the other nsp16_B_ via electrostatic and van der Waals interactions (**Fig. 2F**). Moreover, the new interface gets additional support from the nsp10-nsp10 interface. A residue pair of Thr49 and Met63 from opposing nsp10 protomers mediates hydrophobic and water-mediated interactions. Altogether, Thr49-Met63 pairs add BSA of ∼ 85.00 Å^2^ of BSA to the heterotetrametric interface (**Fig. 2E**).

### Extended RNA-binding groove and regulation of 2’-*O*-methylation

The two gate loops in nsp16 surround the terminal portion of the cap (akin to previously reported structures)^8, 12^ and target A_1,_ whereas the U_2_ stacks between A_1_ purine and the side chain of Asp75. Their Watson-Crick-Franklin edges face the solvent. The U_3_ emerging from this nsp16_A_ takes a sharp 180° turn to face the gate loop 1 of nsp16_A_ and the N-terminal η1 helix of nsp16_B_. The base of U_3_ is held in place by direct h-bonds with the backbone of Ser33 and Ala34, whereas its ribose is stabilized by the Ser33 side chain and coordinating waters with a magnesium ion (**Fig. 2F, S1E**). During the pandemic, a SARS-CoV-2 variant that circulated in New York City harbors Ser33Arg mutation in nsp16^28^. The loss in enzymatic activity when Ser33 was mutated to Arg^12^ or Ala now explains the significance of Ser33 in a clinical variant (**Fig. 2G**). The N-helix of nsp10_A_ faces the bound RNA, which is in close proximity with a distance <13 Å, thus revealing the RNA binding capability of nsp10. Consistently, the truncated nsp10 lacking the N-helix showed a marked reduction in RNA methylation (**Fig. 2G**).

Interestingly, the inverted arrangement of the two nsp16 dimers creates a continuum of positively charged surfaces that extends from the catalytic surface of one of the protomers (nsp16_B_) to the back side of the other protomer (nsp16_A_), suggesting that the downstream portion of the target mRNA emerging from the catalytic region of one nsp16 could potentially be interacting with the positively charged surface on the backside of the other nsp16 (**Fig. 2H**). The tetramerization of nsp16/nsp10 creates a continuum of positively charged surfaces downstream of the target adenine (A_1_) to which the extended RNA beyond U_3_ may bind (**Fig. 2H**). An obvious question arises as to why we did not observe strong electron density for RNA bases beyond U_3_. There could be several explanations for it: a) the broad distribution of positively charged surface is optimal for binding to a complete stem-loop sequence (SL1, **Fig. 1B**) that is lacking in the 10-mer Cap-0 RNA we used in our crystallographic studies. To this front, we examined various structures of nsp16/nsp10 previously solved in the presence of bound sulfate (SO_4_) ions that mimic the phosphates of RNA (PDB ID: 6WQ3, 6WRZ, 6WVN)^9^. Interestingly, a superposition of these structures onto the nsp16/nsp10 tetramer revealed the positions of sulfate ions (S1 - S10) along the entire region of the newly formed positive surface, suggesting a possible interaction of downstream RNA with this positively charged region (**Fig. 2H**). Since the 10-mer RNA used in co-crystallization only contains a partial sequence of SL1 stem-loop and thus lacks secondary structure, it is conceivable that a linear 10-mer RNA remains flexible in this region of nsp16. b) the complete methylation of A_1_ into A_m1_ occurred *in crystallo*, the product 10-mer Cap-1 RNA could facilitate the exit of RNA from the binding surface, or c) the 3’-end of RNA downstream to target A_1_ base could sample multiple conformations. Thus, a wide RNA binding site may allow the 10-mer RNA to bind at different positions on nsp16 and generate a large B-factor; hence low electron density.

Previously, we and others reported the existence of a distantly located putative ligand/nucleotide-binding pocket in nsp16 at the opposite face of the catalytic pocket^8, 9^. This pocket is constituted by residues from β1/α1 loop (Asn13), C-terminal end of α2 (Thr56, Leu57, Thr58), α7 (Ser276), β8 (Trp189), and β9 (Cys209)^8^ (**Fig. 3A, B**), whereas the N-terminals of α2, α7, β8, and β9 participate in catalysis, RNA recognition, and oligomerization (**Fig. S3A**). Of note, Cys209 was only present in SARS-CoV-2, but not any other member of β-CoVs (**Fig. 3B**). Though this pocket is known to bind a variety of nucleotides and ligands (**Fig. 3A**), we did not observe a clear electron density that can account for any known ligands. Thus, the precise mode of RNA interaction with this region remains unclear. We speculated that the 3’-end of a more extended RNA substrate (>10 nts) may be stabilized by this pocket. Single (Asn13/Leu57/Thr58/Trp189/Cys209), double (Leu57, Trp189), and triple (Leu57, Thr58, Trp189) mutant enzymes with Ala as a replacement for residues that line this pocket showed a reduction in 2’-*O*-ribose methylation, to varying degrees. We mutated Glu173 and Ser202 residues that stabilize the terminal ^m7^G base. We observed a similar decrease in enzymatic activity of the triple (Leu57, Thr58, Cys209) or double (Leu57, Trp189) mutants corresponding to the putative nucleotide-binding pocket (**Fig. 2G**). Interestingly, next to the putative nucleotide-binding pocket across the α2/β2 loop, the two Arg residues (Arg279, on α7/β11 loop, and Arg255, on η5/β10 loop) are spaced by ∼ 8.6Å (Cα - Cα) in Cap-0 analog bound (6WKS), Cap-0 RNA (7JYY) or Cap-1 RNA (7L6R, and the current model) structures. However, their guanidine groups orient differently. For example, these are spaced by ∼ 11.3 Å (C_ζ_ - C_ζ_) in Cap-0 analog bound form (6WKS) but in the Capped-RNA bound structures, their side chains are flipped inward to facilitate unusual interaction to significantly reduce the C_ζ_ - C_ζ_ distance to ∼ 6.0 Å in Cap-0 RNA (7JYY), ∼ 7.3 Å in Cap-1 6-mer RNA (7L6R), ∼ 5.0 Å in Cap-1 10-mer RNA (current study) (**Fig. S3B**). As a result of this radical rearrangement of Arg residues, a strong, positively charged surface is formed near the distant putative nucleotide binding site, which might have a role in stabilizing the 3’-end of a longer RNA. In sum, 2’-*O*-methylation of SARS-CoV-2 mRNA is regulated by the cooperative assembly of nsp16/nsp10 complex mediated by their N-terminal regions, putative nucleotide binding pocket of nsp16, in addition to the catalytic pocket of nsp16 and canonical heterodimeric interface. Thus, the positively charged extended surface created by nsp16 dimers and its putative nucleotide binding pocket modulates the 2’-*O*-MTase activity.

**Fig. 3.**
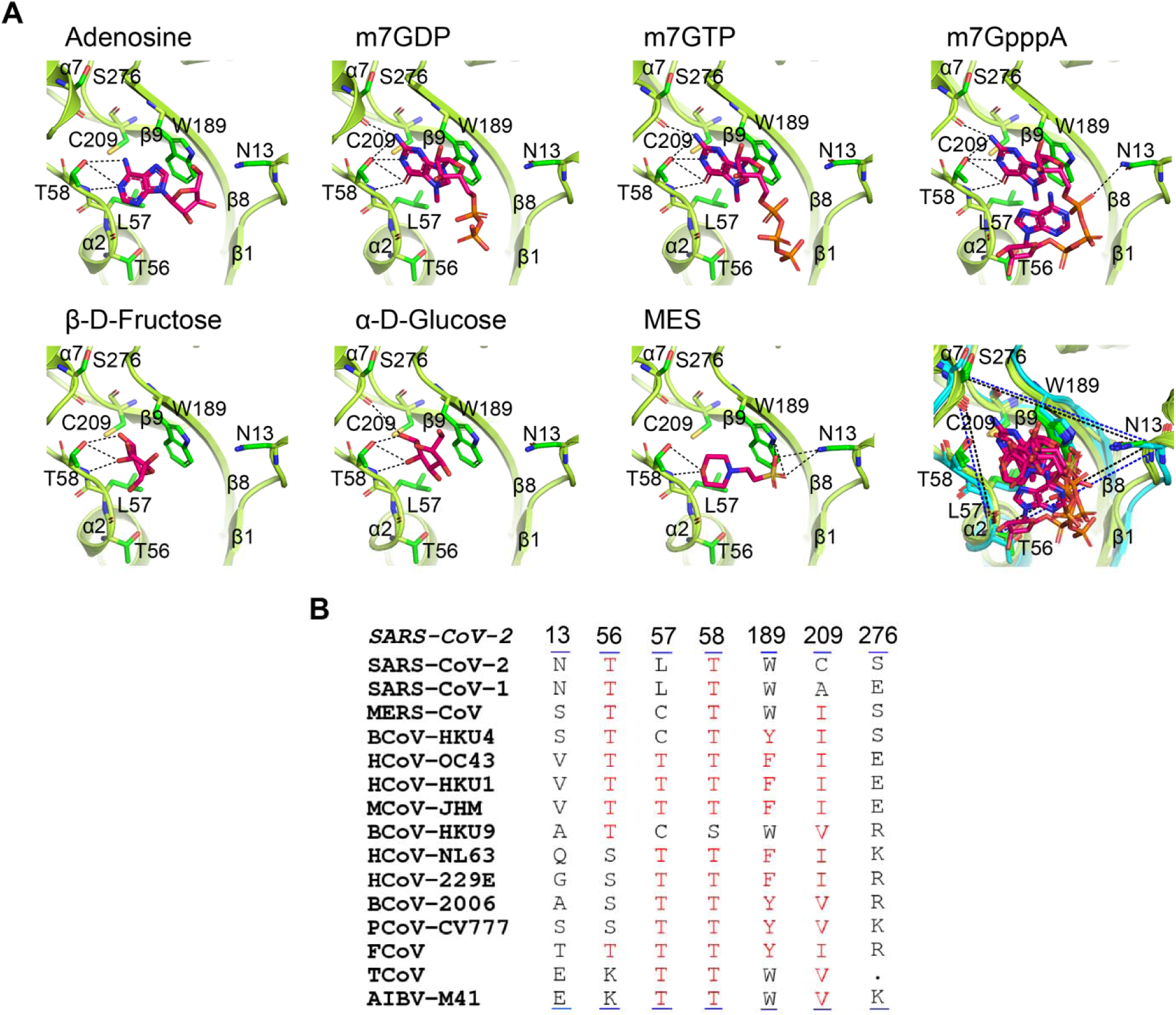
Putative ligand binding pocket of nsp16. **A.** Snapshots of various ligands that bind to the pocket in different nsp16/nsp10 crystal structures: Adenosine (PDB ID: 6WKS), m^7^GDP (PDB ID: 6WQ3), m^7^GTP (PDB ID: 6WVN), m^7^GpppA (PDB ID: 7KOA), β-D-Fructose (PDB ID: 6W4H), α-D-Glucose (PDB ID: 7L6R), MES (PDB ID: 8BSD). Last panel: overlay of ligands in the allosteric pocket represented within a triangular boundary formed by Asn13, Ser276, and Thr56. **B.** Multiple sequence alignment of nsp16 from different CoV members. The alignment for key residues that form the putative ligand-binding pocket in nsp16 and participate in ligand binding is shown (Asn13, Thr56, Leu57, Thr58, Trp189, Cys209, Ser276).

### Requirement of N-terminal domains of nsp10 and nsp16 for RNA stabilization

Nsp10 is an obligate noncatalytic partner of nsp16, but its role in RNA stabilization was unclear. The N-terminal of nsp10 harbors two helices, H1 (N-helix or Helix-1, Ser11 – Phe19) and H2 (Helix-2, Ala23 - Ala32) (**Fig. 4A**). The H2 is conformationally stable and rigid in all the available nsp10 structures whereas H1 (N-helix) is more dynamic and samples different conformations. Many nsp10 structures lack the H1 and preceding N-terminal residues. Previous nsp10 structures in apo and in complex with nsp16 or nsp14 reveal two different orientations of H1 with respect to H2, i.e., a *linear* and a *fold-back* arrangement (**Fig. 4B**). A *linear* arrangement is observed in the presence of nsp16. In contrast, *fold-back* conformation is observed for stand-alone nsp10 or with nsp14. For the first time, the current tetrameric complex co-crystallized with longer RNA reveals a fully structured (up to Val7) N-helix of nsp10 in a *fold-back* conformation in SARS-CoV-2 (**Fig. 4C**). A similar conformation of N-helix was observed in the sinefungin-bound nsp16/nsp10 complex of the OC43-CoV^31^. The stability of nsp10 N-helix in *fold-back* conformation is strengthened by a network of hydrogen bonds between Asn10 and Thr39, which in turn h-bonds (water-mediated) with Asn40, and Arg78 (α3 helix) of nsp10 and backbone carbonyl of Lys76 of nsp16 located at the nsp16-nsp10 canonical interface (**Fig. S3C**). Of note, the side chain of Lys76 h-bonds with the phosphoryl oxygen of the U_3_ base. Pro8 in N-helix, a key residue in SARS-CoV’s nsp10 (**Fig. 4A**), appears to play a critical role as a helix breaker to provide the N-terminal region of nsp10 (Ala1 - Val7) with the flexibility to interact with nsp16-bound RNA. We could only resolve nsp10 N-terminus from Val7 onwards, which is ∼13.0 Å away from PO_4_ of A_4_ base. A similar arrangement of N-helix is observed in the nsp10/nsp14/RNA complex, where it orients the N-terminus to interact with RNA^32^ (**Fig. S3D**). A truncated nsp10 lacking the N-helix (nsp16/^Δ17^nsp10) showed a significant reduction in enzymatic activity (**Fig. 2G**), thus underscoring the critical role of the N-terminal region of nsp10 in efficient 2’-*O*-methylation of SARS-CoV-2 RNA (**Fig. 4D**).

**Fig. 4.**
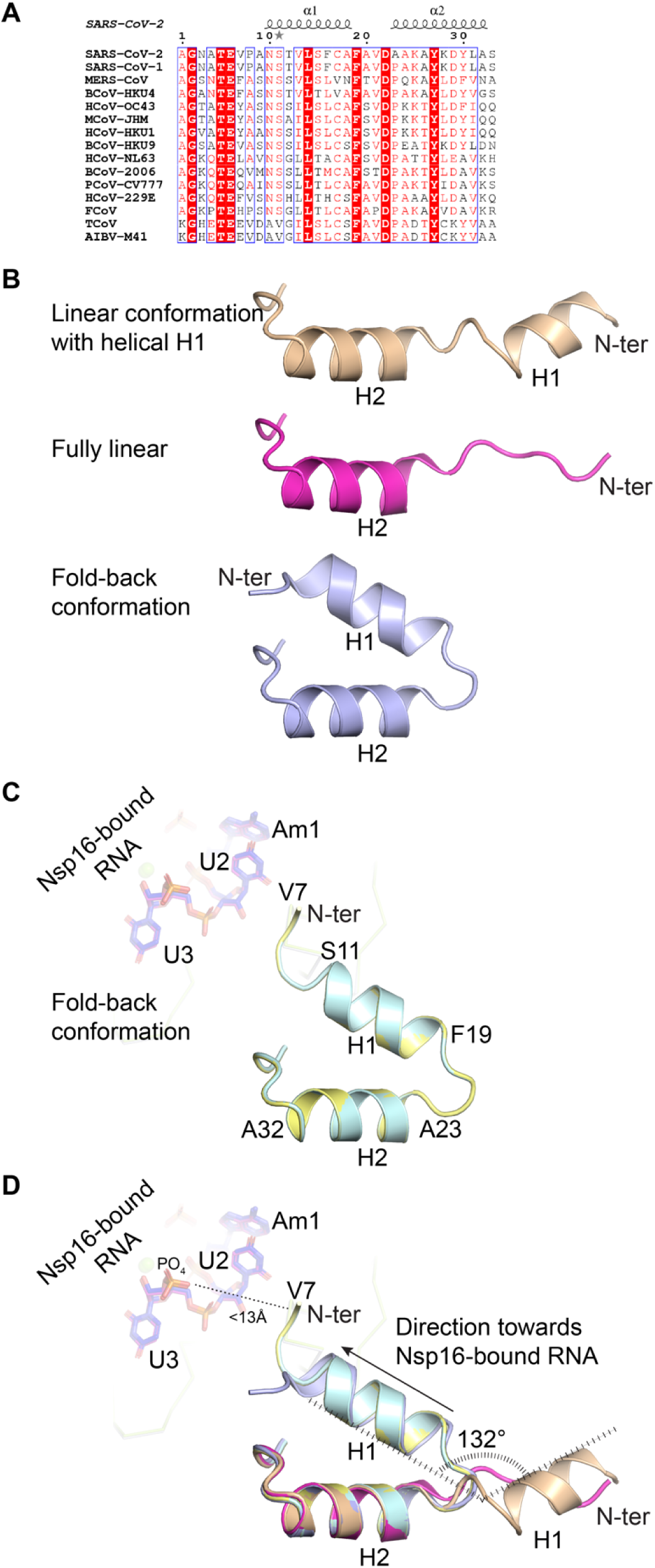
Conformational variability of N-helix of nsp10. **A.** Multiple sequence alignment of the N-terminus of nsp10 from different CoVs. The N-terminal of nsp10 constitutes two major helices, H1 (α1, N10 - A18) and H2 (α2, A23 - A32), and samples two conformations. **B.** linear form displays fully extended and directionally aligned H1 and H2 (PDB ID: 7JYY, chain B and chain D), whereas in a fold-back conformation, H1 folds back on H2 as observed in the apo structure of nsp10 (PDB ID: 6ZCT^36^) or when bound to nsp14/RNA complex (PDB ID: 7N0B). **C.** nsp10 N-terminal prefers a fold-back conformation in the presence of 10-mer RNA (Cap-1) in the current structure; both protomers are shown and overlayed (Cyan: protomer A, Yellow: protomer B), N-helix directing towards the bound RNA of the same protomer in the tetrameric complex. **D.** fold-back conformation superposed onto the linear conformation, depicting the H1 fold-back angle as 132° towards the bound RNA.

A comparative analysis of nsp16 from CoV strains representing three classes (α, β, Y) revealed that in the nsp16 of a CoV strain that infects bats, this aromatic zipper pair (Tyr242, Phe245) is replaced by Asn242 and Leu245. In addition, the first four residues at the N-terminal of nsp16 also show a sequence variation (**Fig. S2**). Only SARS-CoV-2 nsp16 has Ser as the first residue. This Ser is absent, and Lys35 replaces Thr35 in CoVs that infect bats (**Fig. S2)**. The region encompassing Ser2 - Trp5 adopts a 3_10_ helical (η1) conformation, facilitating nsp16-nsp16 dimerization. Notably, the new protein-protein interface we discovered in the tetrameric complex will likely be conserved in most CoV strains but not in those infect bats. Considering our results, it is plausible to speculate that a stable ‘tetrameric complex’ of nsp16/nsp10 might not be formed in CoVs that infect bats. Consequently, it may lead to inefficient or sub-optimal 2’-*O-*methylation (a potent suppressor of innate immunity) and, therefore, a robust innate immune response at a very early stage of infection.

To determine the oligomeric state of nsp16/nsp10/RNA complexes in solution, we probed the complex by size-exclusion chromatography coupled with multi-angle light scattering (SEC-MALS) and analytical ultracentrifugation (AUC). However, the dissociation of nsp16 and nsp10 proteins during both these experiments made the data interpretation inconclusive (data not shown). Therefore, we employed a chemical crosslinking approach followed by immunoblotting of crosslinked species run on SDS-PAGE (**Fig. 5**). We used a BS3-crosslinker to crosslink and capture the oligomeric species of nsp16/nsp10 in solution in the presence and absence of the substrate Cap-0 RNA. To identify proteins in individual bands (bands 1-6), we ran two gels, one used for Coomassie staining and the other for immunoblotting using SARS-CoV-2 nsp16 and nsp10 specific antibodies. Band 1, representing a species of molecular mass of ∼ 14.8 kDa in **Fig. 5A**, was confirmed as nsp10, as visible in the nsp10 immunoblot (**Fig. 5C)**. Band 2 (∼ 33.4 kDa) was confirmed by nsp16 antibody (nsp16 immunoblot, **Fig. 5B**). Band 3, a broader band in the Coomassie-stained gel (**Fig. 5A**), appeared in the presence of the BS3 crosslinker represents a mixture of nsp16/10 (3_A_ in **Fig. 5B, C**) and oligomers of nsp10 (3_B_, and 3_C_ in **Fig. 5C**). Band 4 in **Fig. 5A** represents crosslinked nsp16-nsp16 homodimer which is exclusively detected by nsp16 antibody (**Fig. 5B**); however, this band is missing in the nsp10 immunoblot. This suggests that nsp16 molecules exist in solution close enough to form a homodimer. Band 5 in **Fig. 5A** is detected with both antibodies (**Fig. 5B, C**), which suggests that nsp16 homodimer exists with one molecule of nsp10. Band 6_A/B_ in **Fig. 5A** present in low intensity representing species whose molecular mass corroborates with heterotetrameric (6_A_, ∼ 96.4 kDa) and/or homotrimeric form of nsp16 (6_B_, ∼99.9 kDa). A previous study observed a second nsp16-nsp16 dimeric interface. Thus, the 6_B_ could represent a nsp16 homotrimer with nsp16 crosslinked at both interfaces (**Fig. 6A - C**). Probing with nsp16 and nsp10 antibodies confirms both oligomeric species.

**Fig. 5.**
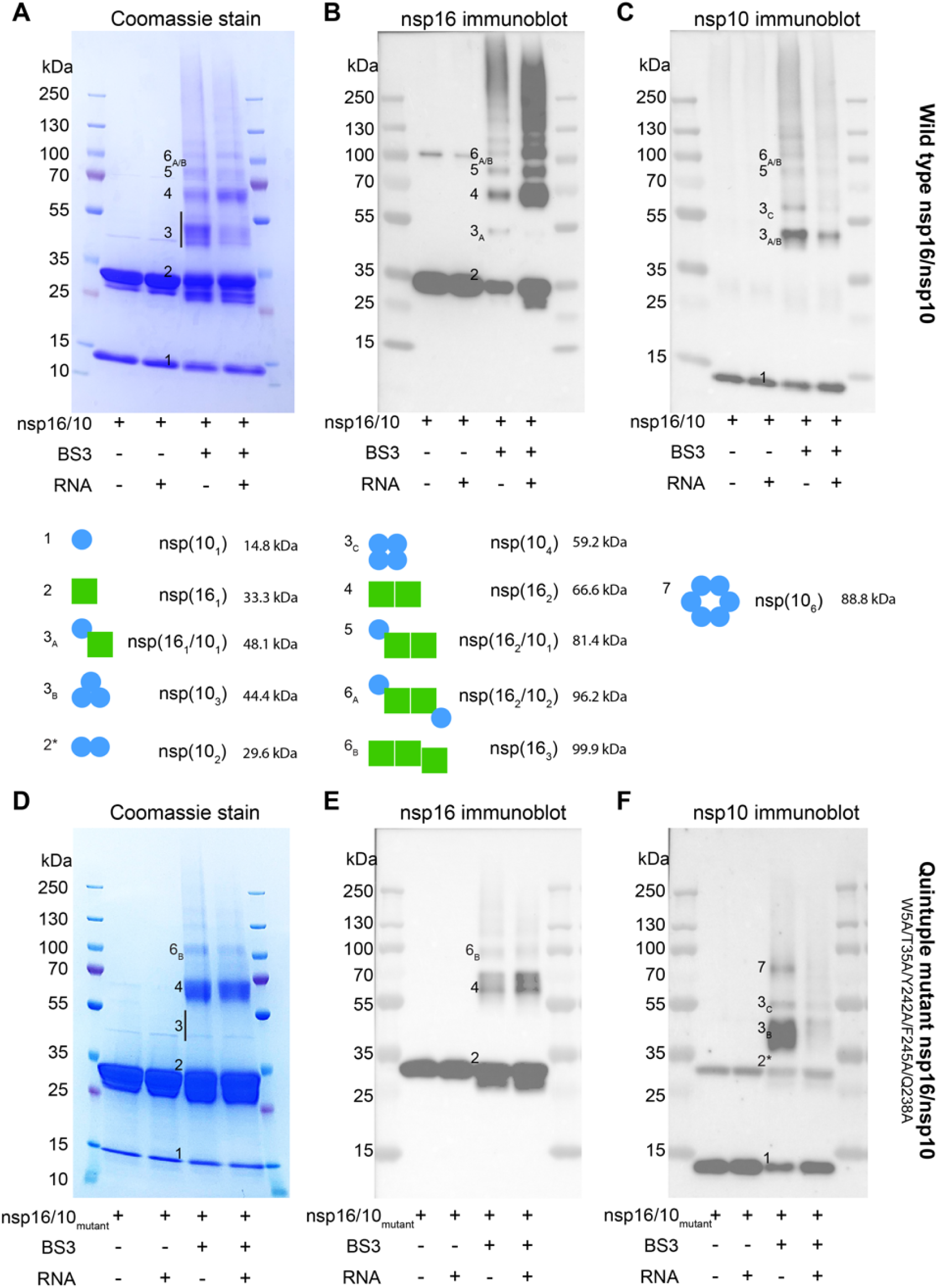
Sampling oligomeric conformations in solution by chemical cross-linking followed by immunoblotting. **A.** Coomassie staining of BS3-crosslinked nsp16/nsp10 (±RNA) on SDS-PAGE. Observed protein bands are labeled as 1-6. **B.** Immunoblotting with nsp16 antibody. Band 2 and 4-6 show the presence of nsp16, whereas part of wider band 3 (3_A_) also represents nsp16. **C.** Immunoblotting with nsp10 antibody. Band 1, 5, and 6 show the presence of nsp10, whereas two parts of wider band 3 (3_B_ and 3_C_) also show nsp10. Cartoon representations of their oligomeric states are in the bottom panel. Blue circle, nsp10; green square, nsp16. **D.** Coomassie staining of BS3-crosslinked quintuple mutant of nsp16/nsp10 (±RNA) on SDS-PAGE. Observed protein bands are labeled as 1-6. **E.** Immunoblotting with nsp16 antibody. **F.** Immunoblotting with nsp10 antibody.

**Fig. 6.**
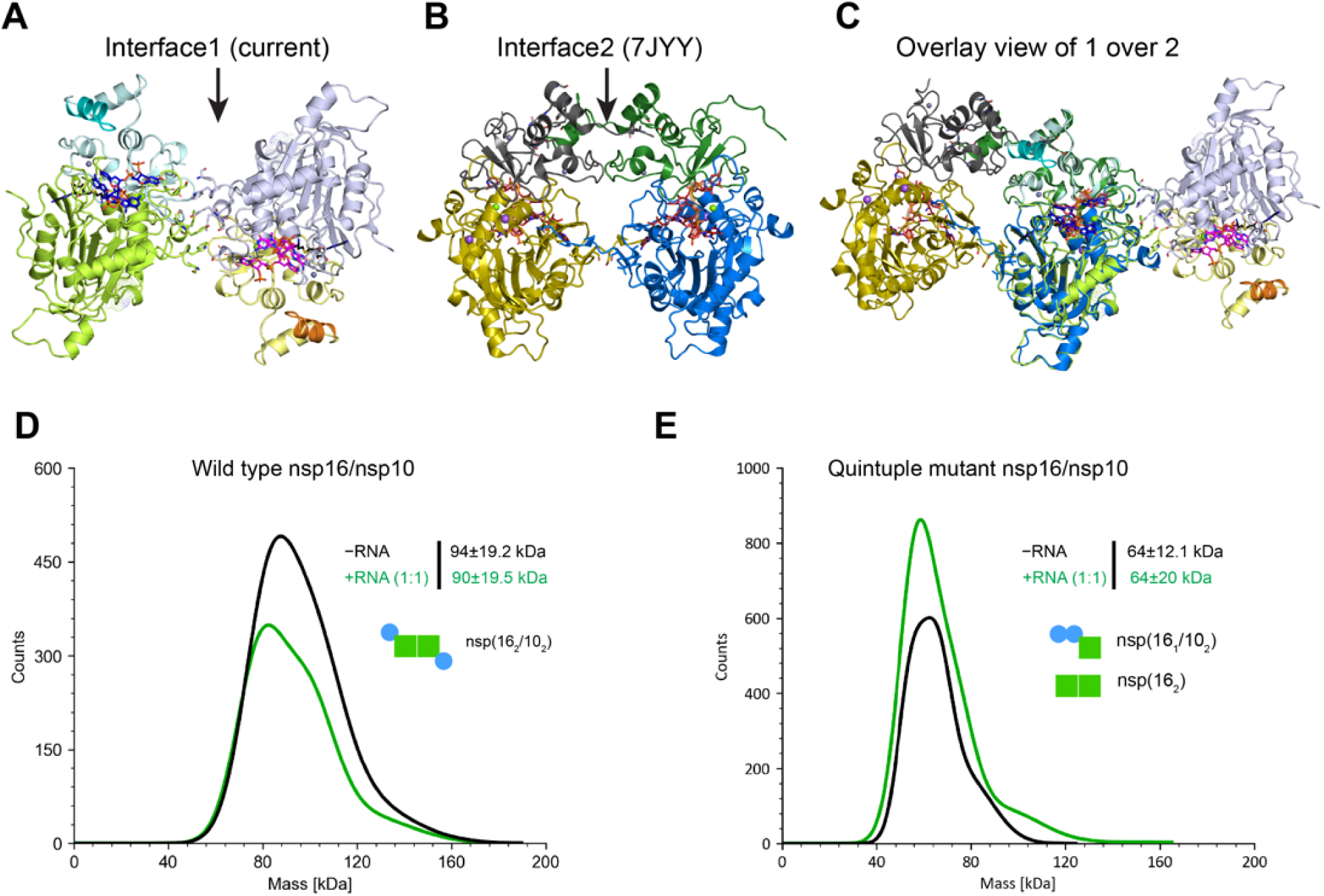
Dimeric interfaces in nsp16/nsp10 complex and investigation of oligomeric conformations using mass photometry. **A.** nsp16-nsp16 interface 1 in the current structure. **B.** nsp16-nsp16 interface 2 in previously reported structure (PDB ID: 7JYY). **C.** An overlay of these two heterotetrameric structures showing two different interfaces on opposite faces of nsp16/nsp10 heterodimer. **D.** Mass photometry analysis of the wild type apo (gray) and with 1:1 molar stoichiometric ratio of RNA (dark green) and quintuple mutant (**E)**.

Interestingly, the increased intensity of band 4 in the presence of RNA suggests a positive effect of RNA in nsp16-nsp16 crosslinking and homodimerization (**Fig. 5A, B**). In contrast, nsp10 undergoes a good amount of self-oligomerization in the absence of RNA, as observed in band 3_B_ (nsp10 trimer) and band 3_C_ (nsp10 tetramer) (**Fig. 5C**). Thus, the crosslinking experiments with immunoblotting confirm the propensity of nsp16/nsp10 to form a heterotetramer in solution. At the same time, the RNA facilitates the formation of the nsp16-nsp16 dimer, the central core of the heterotetrameric nsp16/nsp10 complex. To check the importance of the nsp16-nsp16 interface, we mutated key residues at this interface and purified a quintuple mutant (W5A/T35A/Y242A/F245A/Q238A). A significant loss in RNA methylation activity of this mutant enzyme confirms the functional importance of nsp16 dimerization (**Fig. 2G**). Next, we repeated the crosslinking experiments with this mutant enzyme under identical experimental conditions and in the presence or absence of Cap-0 RNA similar to the wild type enzyme (**Fig. 5D - F**). We observed some additional interesting results with quintuple mutant. Similar to the wild type, the quintuple mutant also showed band 1 (∼ 14.8k Da, nsp10) (**Fig. 5D, F**). A more intense band 2 in Fig. 5D was confirmed as a mixture of nsp16 (major proportion) and nsp10 (minor species). Based on the molecular mass, this new species of nsp10, designated as band 2*, would correspond to a nsp10 dimer. Band 2 contains nsp16 as confirmed by the nsp16 antibody (**Fig. 5E**). In contrast to the crosslinking pattern of the wild type enzyme, band 3 is not visible in quintuple mutant’s Coomassie staining or nsp16 immunoblot (**Fig. 5D, E**). However, the nsp10 antibody detected two oligomeric states of nsp10 as 3_B_ (trimer) and 3_C_ (tetramer) nsp10. Band 4 (nsp16-nsp16 dimer) is present in the quintuple mutant, but its intensity in both reactions (+/- RNA) was significantly reduced compared to the wild type enzyme, suggesting the requirement of the aromatic zipper motif for nsp16 dimerization. These mutations, however, could not completely abolish the nsp16-nsp16 dimerization, suggesting an alternate nsp16-nsp16 interface, perhaps the one reported previously such that the mutant for one interface (current study), nsp16 can utilize another interface to still form nsp16-nsp16 dimer (**Fig. 6A – C**).

Similar to the wild type, the mutant enzyme also showed band 6. However, this was confirmed with nsp16 but not the nsp10 antibody. Therefore, band 6 here contains oligomers of nsp16, i.e., a nsp16 trimer (band 6_B_) formed with an unmutated/alternate interface. The absence of nsp16 in band 6 (**Fig, 5F**) indicates the inability of the quintuple mutant to form a heterotetramer. A new band 7, in nsp10 immunoblot (**Fig. 5F**), which was missing in Coomassie and nsp16 immunoblot, may represent a higher oligomer of nsp10 (hexamer). Thus, the wild type enzyme but not its quintuple mutant forms nsp16/nsp10 tetrameric or even higher-order oligomers (compare lanes 4 and 5 in **Fig. 5B and E**).

To determine the molecular mass of nsp16/nsp10/RNA complexes, we conducted mass photometry experiments for the wild type and quintuple mutant (W5A, Y242A, F245A, Q238A, T35A) enzymes (**Fig. 6D, E**). The wild type protein showed a molecular mass of 94 ± 19.2 kDa, corroborating with the calculated mass of a nsp16/nsp10 tetramer (∼ 96.2 kDa) (**Fig. 6D**). However, the molecular mass of the quintuple mutant was observed as 64 ± 12.1 kDa, which is higher than the calculated mass of nsp16/nsp10 heterodimer (∼ 48.1 kDa) (**Fig. 6E**). This mass difference can be attributed to an extra nsp10 associated with the nsp16/nsp10 heterodimer (∼ 62.9 kDa). Such observation is there in **Fig. 5F**, where nsp10 dimer species are detected in the quintuple mutant. Alternatively, the nsp16 quintuple mutant can still dimerize with the other interface (∼ 66.6 kDa), broadly closer to the observed mass. However, no change occurs in their oligomeric states (wild type and mutant) in the presence of the Cap-0 RNA (**Fig. 6D, E**). Thus, the mass photometry data confirm that the wild-type protein, but not the mutant enzyme, can exist as a stable heterotetramer in solution, further supporting our structural and chemical crosslinking data.

## Discussion

The correct chemical makeup and architecture of the 5’-end of viral RNA help SARS-CoV-2 evade the host immune restriction and hijack the host’s protein synthesis machinery for replication. SARS-CoV-2 genome encodes nsp14 and nsp16, two distinct methyltransferases that install a methylation mark on *N*^7^ of terminal guanine of RNA cap and 2’-OH of ribose of the first transcribed or Cap-adjacent nucleotide (A_1_), respectively. The resulting RNA cap mimics the host RNA. It avoids immune surveillance by host sensor MDA5 or IFN-I-stimulated genes IFIT1 and IFIT3 that would otherwise recognize and sequester the uncapped viral RNA and trigger antiviral signaling pathways. We present the first co-crystal structure of the nsp16/nsp10 complex obtained in the presence of a 10-mer Cap-0 RNA containing the cognate SARS-CoV-2 mRNA sequence. Surprisingly, the structure reveals a unique architecture of the two nsp16/nsp10 heterodimers generating a continuum of positively charged surfaces, which a longer RNA could utilize to stabilize 3’-end (**Fig. 1D, 2H**). Such an inverted orientation of nsp16 allows their front surface to engage in the catalytic transfer of methyl group from SAM to the target adenine and back surface, possibly to act in trans to stabilize the downstream RNA emerging from the catalytic center of the opposing nsp16. A putative nucleotide binding pocket on the back surface may stabilize the 3’-end of RNA (**Fig. 2F, 3A, Fig. S3A**). The nucleotide specificity of this pocket remains unclear due to the lack of strong electron density of RNA around this region. The mutagenesis data support its crucial role in the 2’-*O*-MTase activity. Since the SARS-CoV-2 mRNA downstream to the 10-mer RNA used in our crystallography experiments folds into a stem-loop SL1, this pocket could play an architectural role in recognizing the shape of RNA instead of sequence. The formation of a wider positively charged surface by nsp16 dimerization may also facilitate the interaction with host RNAs with dynamic structures such as the splice site of U1 and the branch point site of U2 small nuclear RNAs, to globally suppress the RNA splicing during viral infection^23^. Future studies can uncover the precise mode of nsp16’s interaction with the components of the host splicing machinery.

Another exciting aspect of the tetrameric structure is the proximity of the nsp10 N-helix to RNA, suggesting its importance for enzymatic activity. nsp14/nsp10 can form a tetrameric assembly^31^, and with the nsp16/nsp10 complex also showing the existence of a unique tetrameric structure, the protein oligomerization thus appears to be a critical strategy for SARS-CoV-2 methyltransferases to modify and shield viral RNA from detection by host immune sensors, even during the act of methylation. Our chemical crosslinking experiments, followed by immunoblotting, confirm the presence of tetrameric species in the solution regardless of the presence of Cap-0 RNA. However, the addition of RNA appears to promote an increased formation of the nsp16-nsp16 dimer, which constitutes the core of the heterotetrameric nsp16/nsp10 complex. Additionally, when interface residues were mutated (quintuple mutant in this study), the enzyme could no longer form higher-order nsp16 oligomers involving the nsp16-nsp16 interface reported in this study. Interestingly, the 2’-O methylation activity of VP39, the nsp16 counterpart in vaccinia virus, significantly increased upon extending the length of substrate RNA from 5 to 20 bases, likely due to the stabilization of downstream RNA by a secondary RNA binding site^33^. Thus, a positive effect of longer RNA extends beyond CoVs and could be a universal requirement for efficient 2’-*O* methylation of mRNAs by viral methyltransferases.

One limitation and interpretation of this study is our inability to co-crystallize the nsp16/nsp10 with capped RNA with the entire stem-loop 1 (SL1) region of SARS-CoV-2 mRNA. Future investigations with resolved structures of nsp16/nsp10 with a longer capped RNA encompassing the SL1 should clarify the atomic-level details of protein-RNA interactions surrounding the putative nucleotide-binding site.

## Supporting information

PDB Validation Report

## Acknowledgments

This work was supported by funding from the NIH (R01AI161363 and U19AI171954). Y.K.G. is grateful to the Robert A. Welch Foundation’s support (AQ-2101-20220331). T.V. was supported by the Cancer Prevention and Research Institute of Texas CPRIT (RP170345). R.S.H. is an Investigator of the Howard Hughes Medical Institute, a CPRIT Scholar, and the Ewing Halsell President’s Council Distinguished Chair at the University of Texas Health San Antonio. This work is based on research conducted at the Northeastern Collaborative Access Team beamlines, funded by NIH (P30GM124165) and U.S. Department of Energy (DOE) Contract DE-AC02-06CH11357.

## Author contributions

Y.K.G. conceived, designed, and supervised the study; A.M. performed all crystallographic studies; R.R., S.A., T.V. performed protein purification and biochemical assays; R.R. and M.P. performed chemical cross-linking experiments and immunoblotting with the guidance of L.M.S.; S.Q. assisted with radiometric assays; S.-H.C, N.J., R.S.H, L.M.S. provided reagents. A.M. and Y.K.G. wrote the manuscript with input from all co-authors. All authors read and approved this version.

## Data and materials availability

The information about coding sequences of nsp16 (NCBI reference sequence YP_009725311.1) and nsp10 (NCBI reference sequence: YP_0009725306.1) of the seafood market pneumonia SARS-CoV-2 isolate Wuhan-Hu-1 (NC_045512) used in this study is available at NCBI (https://www.ncbi.nlm.nih.gov/nuccore/NC_045512). The atomic coordinates and structure factors files were deposited in the Protein Data Bank (PDB) under accession code 8VUO. The PDB DOI for this deposition is 10.2210/pdb8vuo/pdb. A PDB validation report is included in supplemental data. All other relevant data are available from the corresponding author via email.

## Methods

### Protein expression and purification

The coding sequences of SARS-CoV-2 nsp16 (NCBI reference sequence YP_009725311.1) and nsp10 (NCBI reference sequence: YP_0009725306.1) of the Wuhan seafood market pneumonia virus isolate Wuhan-Hu-1 (NC_045512) were cloned into a single pETduet-1 vector downstream to a 6xHis-SUMO tag sequence. This plasmid was transformed into an *E. coli* expression strain NiCo21(DE3) (NEB #C2529H) to co-express the nsp16/nsp10 protein complex. The transformed cells were grown in Terrific Broth medium supplemented with ampicillin (100 µg ml^−1^) at 37 °C. Protein expression was induced by adding 0.4 mM isopropyl β-D-1-thiogalactopyranoside (IPTG) at OD_600_ = 0.6-0.8, followed by continued incubation of the cultures for 16 h at 18 °C. Cells from the 2-liter culture were then harvested by centrifugation at 8983 × *g* for 20 min and re-suspended in ice-cold lysis buffer (25 mM Tris-HCl pH 7.5, 0.5 M NaCl, 0.1 mM TCEP, 10% Glycerol, 5mM Imidazole) supplemented with a protease inhibitor tablet (Pierce). Cell lysis was accomplished using a microfluidizer (Analytik, UK), and the soluble fraction was separated by centrifugation at 158,000 × *g* for 40 min. The clarified soluble fraction, after passing through a 0.22 µm filter, was loaded onto a Nuvia IMAC column (Bio-Rad) pre-equilibrated in binding buffer containing 25 mM Tris-HCl pH 7.5, 0.5 M NaCl, 0.1 mM TCEP, 10% Glycerol, 5mM Imidazole. The proteins were eluted by increasing the imidazole concentration from 0 M to 1.0 M). The 6xHis-SUMO tag was then proteolytically removed, and the tag-free sample was re-applied to the IMAC column to separate the uncleaved protein fraction. The proteins were finally purified by passing through the HiLoad 16/600 Superdex 75 column (GE Healthcare). The nsp16/nsp10 complex was eluted as a single homogenous species in a final buffer containing 25 mM Tris-HCl pH 7.5, 0.2 M NaCl, 0.1 mM TCEP, and 5 mM MgSO_4_. The purified protein complex was concentrated to 6 mg/mL, flash-frozen in liquid nitrogen, and stored at −80 °C. We used the same method for all mutant nsp16/nsp10 enzymes reported in this study. We introduced each single point (N13A, L57A, T58A, W189A, C209A), double (L57A/W189A), triple (L57A/T58A/W189 and W5A/Y242A/F245A), and quintuple (W5A/T35A/Y242A/F245A/Q238A) mutations and an N-terminal deletion mutant of nsp10 (nsp16/^Δ17^nsp10) in the nsp16/nsp10 plasmid.

### Crystallization, X-ray data collection, and structure determination

The purified nsp16/nsp10 protein complex (6 mg/mL), mixed with 1.25 molar excess of Cap-0 RNA (^m7^GpppAUUAAAGGUU, dissolved in a buffer containing 10 mM Bis-Tris pH 6.5, 50 mM NaCl, and heated at 80 °C for 5 minutes and then gradually cooled to room temperature overnight). This mixture was subjected to extensive crystallization trials using commercial screens. The ternary complex of nsp16/nsp10/RNA was co-crystallized by the sitting drop vapor diffusion method at 4 °C. The plate-shaped crystals appeared in a crystallization solution containing 25% (v/v) Ethylene glycol in five weeks. Crystals were flash-frozen directly from the original drop into liquid nitrogen without further cryoprotection. The best nsp16/nsp10/RNA complex co-crystal diffracted X-rays to ∼ 2.39 Å resolution with synchrotron radiation (**Table 1**). The crystals belong to space group P2_1_ with unit cell dimensions a = 111.75 Å, b = 58.89 Å, c = 117.02 Å, α = γ = 90°, and β = 95.48°, and with two nsp16/nsp10 heterodimer per asymmetric unit. The X-ray diffraction data measured at wavelength 0.9792 Å were indexed, integrated, and scaled using XDS, aimless, and various ccp4 suite programs (truncate, freeflag, and mtz2various) incorporated into the RAPD pipeline at the NECAT 24ID beamline^34^. The structure was solved by molecular replacement using an nsp16/nsp10 structure bound to SAM and Cap-0 analog (PDB ID: 6WKS) as a template in Phaser. The resulting maps indicated clear electron densities for two nsp16, and two nsp10 protomers, with SAH occupying the catalytic pocket of nsp16. Three of the ten nucleotides, including the RNA cap of the 10-mer RNA (^m7^GpppA_1_•U_2_•U_3_), showed clear electron density. Only the phosphate from the A_4_ nucleotide could also be built with clear electron density. Interestingly, the ribose sugar in the A_1_ nucleotide is methylated at the 2’-*O* position (**Fig. S1B**), attributing to *in crystallo* reaction that results in the complex in product (Cap-1) bound form. We iteratively built and refined the model with good stereochemistry using Coot and REFMAC/Phenix.refine, respectively (**Table 1**). The final model was refined to 2.39 Å resolution with R_free_ and R_work_ values of ∼ 21.86% and ∼ 18.00%, respectively. We generated ligand topologies and geometrical restraints using PRODRG (http://prodrg1.dyndns.org), GRADE (http://grade.globalphasing.org), and eLBOW (Phenix)^35^. All figures of structural models were generated using Pymol (The PyMOL Molecular Graphics System, Version 2.5.4 Schrödinger, LLC). The final figures were prepared using Adobe Illustrator (version 2023).

### Enzyme activity assay

To assess the preference of RNA substrates for nsp16/nsp10, a radiometric assay was performed to test methyltransferase activity for each RNA. Three RNA substrates: Cap-0 10-mer (^m7^GpppAUUAAAGGUU); Cap-0 16-mer (^m7^GpppAUUAAAGGUUUAUACC); Cap-0 33-mer (^m7^GpppAUUAAAGGUUUAUACCUUCCCAGGUAACAAACC), and an RNA Cap-0 analog (^m7^GpppA) were tested (**Fig. 1B**). Each reaction was carried out in a 5 µL mixture containing 50 mM Tris pH 8.0, 5 mM KCl, 1 mM dithiothreitol (DTT), 1mM MgCl_2_, 5 µM [methyl-^3^H] SAM (PerkinElmer), 10 µM substrate RNA, and 2 µM purified nsp16/nsp10 enzyme complex. The reactions were incubated at 37 °C for 1 h, and 4 µL of each reaction was quenched by blotting on the Hybond-N+ membrane (Amersham). RNA probes were crosslinked to the membrane by exposure to ultraviolet light (254 nm) for 2 min. The membranes were successively washed three times with 1X PBS followed by three ethanol washes for 5 min each. The membranes were air-dried in the hood for 15 min, and the counts per minute (c.p.m.) of the RNA probes were measured using a scintillation counter (Beckman LS6500). Results (**Fig. 1C, 2G**) presented here are an average of three independent experiments (n = 3) with standard deviation (shown as error bars) for respective RNA substrates.

### Chemical cross-linking

For the primary amine-based crosslinking experiment, nsp16/nsp10 protein complex was dialyzed in non-amine buffer (20 mM HEPES pH7.5, 200 mM NaCl, 5% Glycerol, 0.1 mM TCEP, 5mM MgSO4). Crosslinking reactions for nsp16/nsp10 (10µM) were done with 0.1 mM bis(sulfosuccinimidyl) suberate (BS3, Thermo Scientific cat no. A39266) in the presence and absence of RNA substrate 16mer Cap-0 RNA (12.5µM) with a final reaction volume of 20 µl. Reactions were incubated for 1 hour at room temperature and then terminated by Tris buffer pH 8.0 at a final concentration of 20 mM. The cross-linked products and uncrosslinked nsp16/nsp10 (control) were analyzed on 4-20% Tris-Glycine SDS PAGE gels **(Fig. 5A, D)**. The gels were stained with Coomassie and imaged on a Biorad ChemiDoc imager.

### Western blotting

BS3 Cross-linked nsp16/nsp10 (±RNA) was quantified using Protein Assay Dye (Bio-Rad, 5000006), denatured, and loaded on tris-glycine SDS PAGE gels (500ng protein in each lane) to separate them. The protein bands were then transferred onto the nitrocellulose membrane (Cytiva, 10600002) and probed using listed primary and secondary antibodies. The signal was detected using Clarity and Clarity Max ECL substrates (Biorad, 1705060, 1705062) and imaged on a Biorad ChemiDoc imager (**Fig. 5B, C, E, F**). The following antibodies were used: SARS-CoV-2 nsp10 (ProSci, 9179), SARS-CoV-2 nsp16 (ProSci, 9271), anti-rabbit IgG HRP secondary (Cell Signaling, 7074).

### Mass photometry

The mass photometry experiments were performed using a TwoMP mass photometer (Refeyn) instrument in a buffer containing 20 mM Bis-Tris pH 6.5 and 150 mM NaCl. Data were acquired using Refeyn AcquireMP software and analyzed by DiscoverMP, v2.3. The molecular mass standard was used as a reference to estimate the molecular mass of nsp16/nsp10/RNA complexes. The mass photometry experiments for the wild-type and quintuple mutant enzyme (W5A, Y242A, F245A, Q238A, T35A), which showed minimal methylation activity (**Fig. 2G**), were conducted at 100 nM concentration in the presence and absence of a 16-mer Cap-0 RNA at a 1:1 molar stoichiometry (**Fig. 6D, E**).

## References

[1] Daffis, S., Szretter, K. J., Schriewer, J., Li, J., Youn, S., Errett, J., Lin, T. Y., Schneller, S., Zust, R., Dong, H., Thiel, V., Sen, G. C., Fensterl, V., Klimstra, W. B., Pierson, T. C., Buller, R. M., Gale, M., Jr., Shi, P. Y., and Diamond, M. S. (2010) 2’-O methylation of the viral mRNA cap evades host restriction by IFIT family members, Nature 468, 452–456.

[2] Ramanathan, A., Robb, G. B., and Chan, S. H. (2016) mRNA capping: biological functions and applications, Nucleic Acids Res 44, 7511–7526.

[3] Nencka, R., Silhan, J., Klima, M., Otava, T., Kocek, H., Krafcikova, P., and Boura, E. (2022) Coronaviral RNA-methyltransferases: function, structure and inhibition, Nucleic Acids Res 50, 635–650.

[4] Park, G. J., Osinski, A., Hernandez, G., Eitson, J. L., Majumdar, A., Tonelli, M., Henzler-Wildman, K., Pawlowski, K., Chen, Z., Li, Y., Schoggins, J. W., and Tagliabracci, V. S. (2022) The mechanism of RNA capping by SARS-CoV-2, Nature 609, 793–800.

[5] Bouvet, M., Debarnot, C., Imbert, I., Selisko, B., Snijder, E. J., Canard, B., and Decroly, E. (2010) In vitro reconstitution of SARS-coronavirus mRNA cap methylation, PLoS Pathog 6, e1000863.

[6] Chen, Y., Su, C., Ke, M., Jin, X., Xu, L., Zhang, Z., Wu, A., Sun, Y., Yang, Z., Tien, P., Ahola, T., Liang, Y., Liu, X., and Guo, D. (2011) Biochemical and structural insights into the mechanisms of SARS coronavirus RNA ribose 2’-O-methylation by nsp16/nsp10 protein complex, PLoS Pathog 7, e1002294.

[7] Decroly, E., Debarnot, C., Ferron, F., Bouvet, M., Coutard, B., Imbert, I., Gluais, L., Papageorgiou, N., Sharff, A., Bricogne, G., Ortiz-Lombardia, M., Lescar, J., and Canard, B. (2011) Crystal structure and functional analysis of the SARS-coronavirus RNA cap 2’-O-methyltransferase nsp10/nsp16 complex, PLoS Pathog 7, e1002059.

[8] Viswanathan, T., Arya, S., Chan, S. H., Qi, S., Dai, N., Misra, A., Park, J. G., Oladunni, F., Kovalskyy, D., Hromas, R. A., Martinez-Sobrido, L., and Gupta, Y. K. (2020) Structural basis of RNA cap modification by SARS-CoV-2, Nat Commun 11, 3718.

[9] Rosas-Lemus, M., Minasov, G., Shuvalova, L., Inniss, N. L., Kiryukhina, O., Brunzelle, J., and Satchell, K. J. F. (2020) High-resolution structures of the SARS-CoV-2 2’-O-methyltransferase reveal strategies for structure-based inhibitor design, Sci Signal 13.

[10] Wilamowski, M., Sherrell, D. A., Minasov, G., Kim, Y., Shuvalova, L., Lavens, A., Chard, R., Maltseva, N., Jedrzejczak, R., Rosas-Lemus, M., Saint, N., Foster, I. T., Michalska, K., Satchell, K. J. F., and Joachimiak, A. (2021) 2’-O methylation of RNA cap in SARS-CoV-2 captured by serial crystallography, Proc Natl Acad Sci U S A 118.

[11] Krafcikova, P., Silhan, J., Nencka, R., and Boura, E. (2020) Structural analysis of the SARS-CoV-2 methyltransferase complex involved in RNA cap creation bound to sinefungin, Nat Commun 11, 3717.

[12] Viswanathan, T., Misra, A., Chan, S. H., Qi, S., Dai, N., Arya, S., Martinez-Sobrido, L., and Gupta, Y. K. (2021) A metal ion orients SARS-CoV-2 mRNA to ensure accurate 2’-O methylation of its first nucleotide, Nat Commun 12, 3287.

[13] Minasov, G., Rosas-Lemus, M., Shuvalova, L., Inniss, N. L., Brunzelle, J. S., Daczkowski, C. M., Hoover, P., Mesecar, A. D., and Satchell, K. J. F. (2021) Mn(2+) coordinates Cap-0-RNA to align substrates for efficient 2’-O-methyl transfer by SARS-CoV-2 nsp16, Sci Signal 14.

[14] Russ, A., Wittmann, S., Tsukamoto, Y., Herrmann, A., Deutschmann, J., Lagisquet, J., Ensser, A., Kato, H., and Gramberg, T. (2022) Nsp16 shields SARS-CoV-2 from efficient MDA5 sensing and IFIT1-mediated restriction, EMBO Rep, e55648.

[15] Zust, R., Cervantes-Barragan, L., Habjan, M., Maier, R., Neuman, B. W., Ziebuhr, J., Szretter, K. J., Baker, S. C., Barchet, W., Diamond, M. S., Siddell, S. G., Ludewig, B., and Thiel, V. (2011) Ribose 2’-O-methylation provides a molecular signature for the distinction of self and non-self mRNA dependent on the RNA sensor Mda5, Nat Immunol 12, 137–143.

[16] Schindewolf, C., Lokugamage, K., Vu, M. N., Johnson, B. A., Scharton, D., Plante, J. A., Kalveram, B., Crocquet-Valdes, P. A., Sotcheff, S., Jaworski, E., Alvarado, R. E., Debbink, K., Daugherty, M. D., Weaver, S. C., Routh, A. L., Walker, D. H., Plante, K. S., and Menachery, V. D. (2023) SARS-CoV-2 Uses Nonstructural Protein 16 To Evade Restriction by IFIT1 and IFIT3, J Virol 97, e0153222.

[17] Schindewolf, C., and Menachery, V. D. (2023) Coronavirus 2’-O-methyltransferase: A promising therapeutic target, Virus Res 336, 199211.

[18] Park, A., and Iwasaki, A. (2020) Type I and Type III Interferons - Induction, Signaling, Evasion, and Application to Combat COVID-19, Cell Host Microbe 27, 870-878.

[19] Almazan, F., Dediego, M. L., Galan, C., Escors, D., Alvarez, E., Ortego, J., Sola, I., Zuniga, S., Alonso, S., Moreno, J. L., Nogales, A., Capiscol, C., and Enjuanes, L. (2006) Construction of a severe acute respiratory syndrome coronavirus infectious cDNA clone and a replicon to study coronavirus RNA synthesis, J Virol 80, 10900–10906.

[20] Menachery, V. D., Yount, B. L., Jr., Josset, L., Gralinski, L. E., Scobey, T., Agnihothram, S., Katze, M. G., and Baric, R. S. (2014) Attenuation and restoration of severe acute respiratory syndrome coronavirus mutant lacking 2’-o-methyltransferase activity, J Virol 88, 4251–4264.

[21] Tsukamoto, Y., Igarashi, M., and Kato, H. (2023) Targeting cap1 RNA methyltransferases as an antiviral strategy, Cell Chem Biol.

[22] Wang, Y., Sun, Y., Wu, A., Xu, S., Pan, R., Zeng, C., Jin, X., Ge, X., Shi, Z., Ahola, T., Chen, Y., and Guo, D. (2015) Coronavirus nsp10/nsp16 Methyltransferase Can Be Targeted by nsp10-Derived Peptide In Vitro and In Vivo To Reduce Replication and Pathogenesis, J Virol 89, 8416–8427.

[23] Banerjee, A. K., Blanco, M. R., Bruce, E. A., Honson, D. D., Chen, L. M., Chow, A., Bhat, P., Ollikainen, N., Quinodoz, S. A., Loney, C., Thai, J., Miller, Z. D., Lin, A. E., Schmidt, M. M., Stewart, D. G., Goldfarb, D., De Lorenzo, G., Rihn, S. J., Voorhees, R. M., Botten, J. W., Majumdar, D., and Guttman, M. (2020) SARS-CoV-2 Disrupts Splicing, Translation, and Protein Trafficking to Suppress Host Defenses, Cell 183, 1325–1339 e1321.

[24] Rangan, R., Zheludev, I. N., Hagey, R. J., Pham, E. A., Wayment-Steele, H. K., Glenn, J. S., and Das, R. (2020) RNA genome conservation and secondary structure in SARS-CoV-2 and SARS-related viruses: a first look, RNA 26, 937–959.

[25] Sola, I., Almazan, F., Zuniga, S., and Enjuanes, L. (2015) Continuous and Discontinuous RNA Synthesis in Coronaviruses, Annu Rev Virol 2, 265–288.

[26] Yang, D., and Leibowitz, J. L. (2015) The structure and functions of coronavirus genomic 3’ and 5’ ends, Virus Res 206, 120–133.

[27] Hodel, A. E., Gershon, P. D., Shi, X., and Quiocho, F. A. (1996) The 1.85 A structure of vaccinia protein VP39: a bifunctional enzyme that participates in the modification of both mRNA ends, Cell 85, 247–256.

[28] Butler, D. J., Mozsary, C., Meydan, C., Danko, D., Foox, J., Rosiene, J., Shaiber, A., Afshinnekoo, E., MacKay, M., Sedlazeck, F. J., Ivanov, N. A., Sierra, M., Pohle, D., Zietz, M., Gisladottir, U., Ramlall, V., Westover, C. D., Ryon, K., Young, B., Bhattacharya, C., Ruggiero, P., Langhorst, B. W., Tanner, N., Gawrys, J., Meleshko, D., Xu, D., Steel, P. A. D., Shemesh, A. J., Xiang, J., Thierry-Mieg, J., Thierry-Mieg, D., Schwartz, R. E., Iftner, A., Bezdan, D., Sipley, J., Cong, L., Craney, A., Velu, P., Melnick, A. M., Hajirasouliha, I., Horner, S. M., Iftner, T., Salvatore, M., Loda, M., Westblade, L. F., Cushing, M., Levy, S., Wu, S., Tatonetti, N., Imielinski, M., Rennert, H., and Mason, C. E. (2020) Shotgun Transcriptome and Isothermal Profiling of SARS-CoV-2 Infection Reveals Unique Host Responses, Viral Diversification, and Drug Interactions, bioRxiv.

[29] Brass, A., Kadler, K. E., Thomas, J. T., Grant, M. E., and Boot-Handford, R. P. (1991) The aromatic zipper: a model for the initial trimerization event in collagen folding, Biochem Soc Trans 19, 365S.

[30] Nair, R. V., Kheria, S., Rayavarapu, S., Kotmale, A. S., Jagadeesh, B., Gonnade, R. G., Puranik, V. G., Rajamohanan, P. R., and Sanjayan, G. J. (2013) A synthetic zipper peptide motif orchestrated via co-operative interplay of hydrogen bonding, aromatic stacking, and backbone chirality, J Am Chem Soc 135, 11477–11480.

[31] Dostalik, P., Krafcikova, P., Silhan, J., Kozic, J., Chalupska, D., Chalupsky, K., and Boura, E. (2021) Structural Analysis of the OC43 Coronavirus 2’-O-RNA Methyltransferase, J Virol 95, e0046321.

[32] Liu, C., Shi, W., Becker, S. T., Schatz, D. G., Liu, B., and Yang, Y. (2021) Structural basis of mismatch recognition by a SARS-CoV-2 proofreading enzyme, Science 373, 1142–1146.

[33] Lockless, S. W., Cheng, H. T., Hodel, A. E., Quiocho, F. A., and Gershon, P. D. (1998) Recognition of capped RNA substrates by VP39, the vaccinia virus-encoded mRNA cap-specific 2’-O-methyltransferase, Biochemistry 37, 8564–8574.

[34] Collaborative Computational Project, N. (1994) The CCP4 suite: programs for protein crystallography, Acta Crystallogr D Biol Crystallogr 50, 760–763.

[35] Adams, P. D., Afonine, P. V., Bunkoczi, G., Chen, V. B., Davis, I. W., Echols, N., Headd, J. J., Hung, L. W., Kapral, G. J., Grosse-Kunstleve, R. W., McCoy, A. J., Moriarty, N. W., Oeffner, R., Read, R. J., Richardson, D. C., Richardson, J. S., Terwilliger, T. C., and Zwart, P. H. (2010) PHENIX: a comprehensive Python-based system for macromolecular structure solution, Acta Crystallogr D Biol Crystallogr 66, 213–221.

[36] Rogstam, A., Nyblom, M., Christensen, S., Sele, C., Talibov, V. O., Lindvall, T., Rasmussen, A. A., Andre, I., Fisher, Z., Knecht, W., and Kozielski, F. (2020) Crystal Structure of Non-Structural Protein 10 from Severe Acute Respiratory Syndrome Coronavirus-2, Int J Mol Sci 21.

