## Supplemental Figures 1-3 for "Structural Insights into the Assembly and Regulation of 2’-*O* RNA Methylation by SARS-CoV-2 nsp16/nsp10"

This supplementary material document contains the following:

Fig. S1-S3

**Fig. S1**

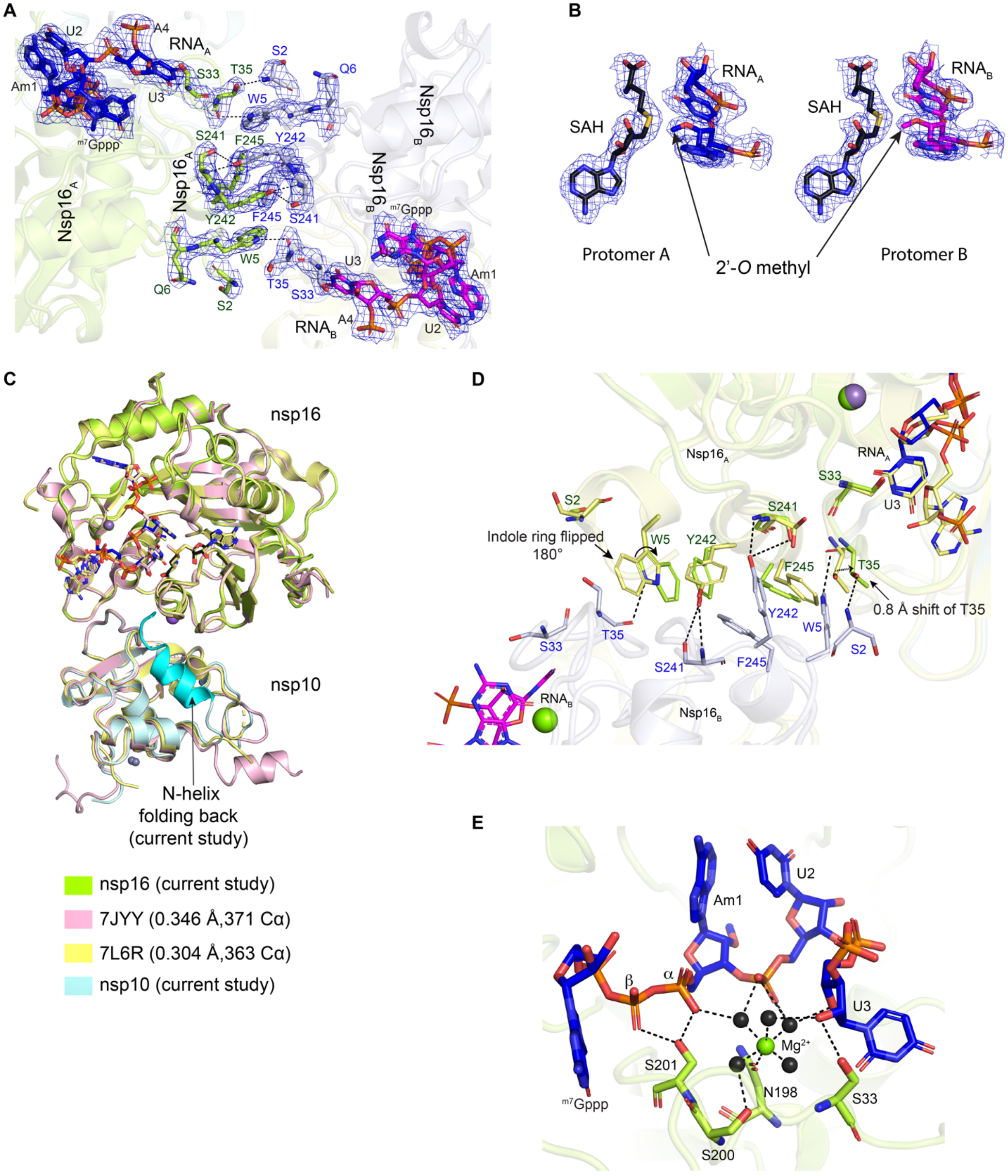

A
RNAA
U2
A4
Am1
U3
m7Gppp
S33
T35
S2
Q6
W5
S241
F245
Y242
Nsp16A
Nsp16B
RNAB
B
SAH
RNAA
RNAB
Protomer A
Protomer B
2'-O methyl
C
nsp16
nsp10
N-helix folding back (current study)
nsp16 (current study)
7JYY (0.346 Å,371 Cα)
7L6R (0.304 Å,363 Cα)
nsp10 (current study)
D
Indole ring flipped 180°
0.8 Å shift of T35
E
β
α
Mg2+
S201
S200
N198

**Supplementary Fig. 1. A.** Two RNA chains running across the aromatic zipper interface of nsp16. The portion of RNA ($^{m7}$GpppAm1U2U3A4$^{PO4}$) refined with occupancy 1.0 could be observed as shown in dark colors (blue and magenta sticks) with 2Fo-Fc map contoured at $\sigma = 1.0$ (blue mesh). **B.** Electron density map for SAH and 2'-*O* methyl group at the A1 nucleotide (2Fo-Fc map contoured at $\sigma = 1.0$) suggesting *in crystallo* methyltransferase reaction occurred for both copies of the nsp16/nsp10/RNA complex. **C.** Structural Comparison of current nsp16/nsp10/RNA complex with the known structures (Cap-0 6-mer RNA: 7JYY, and Cap-1 6-mer RNA: 7L6R). **D.** Indole ring of Trp5 at the N-terminal of nsp16 undergoes a 180° flip to form aromatic zipper at the nsp16-nsp16 interface and Thr35 moves inwards (~ 0.8 Å in comparison to cap-1 bound 6-mer RNA) towards bound RNA to form interaction with terminal Serine (S2) of opposite nsp16 molecule. **E.** Magnesium at the protein RNA interface with Ser33 holding the sugar of $U_3$ while Ser201 stabilizes the α and β phosphoryl oxygens of RNA cap. Green sphere, $Mg^{2+}$ ion; Black sphere, water; Black dashed lines, h-bonds.

**Fig. S2**

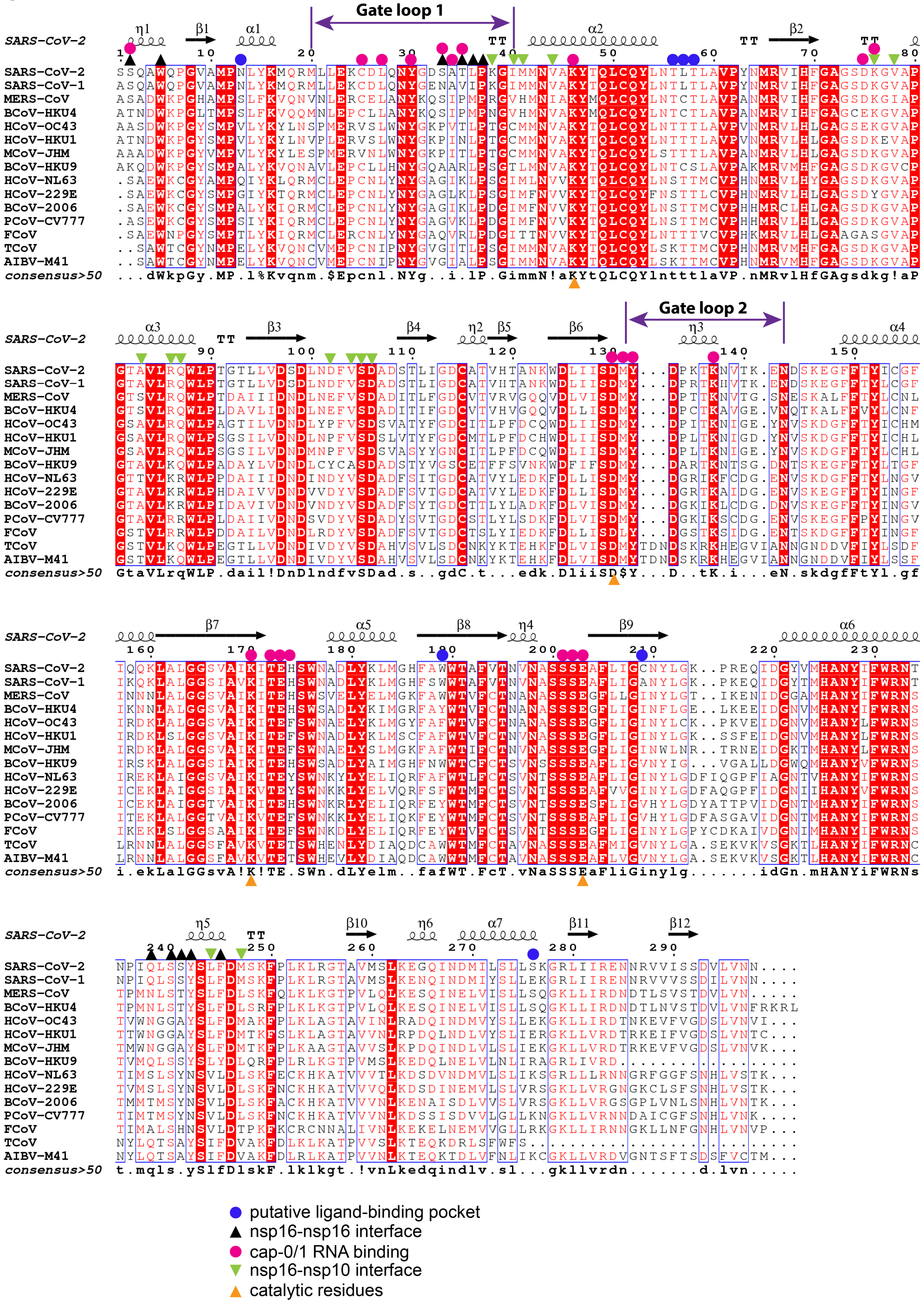
Gate loop 1
Gate loop 2
SARS-CoV-2
SARS-CoV-1
MERS-CoV
BCoV-HKU4
HCoV-OC43
HCoV-HKU1
MCoV-JHM
BCoV-HKU9
HCoV-NL63
HCoV-229E
BCoV-2006
PCoV-CV777
FCoV
TCoV
AIBV-M41
consensus>50
putative ligand-binding pocket
nsp16-nsp16 interface
cap-0/1 RNA binding
nsp16-nsp10 interface
catalytic residues

**Supplementary Fig. 2. Multiple sequence alignment of the protein sequence of nsp16 from 15 CoV members**. (SARS-CoV-2, Uniprot ID: P0DTD1; SARS-CoV, Uniprot ID: P0C6X7; MERS-CoV, Uniprot ID: K9N7C7; BCoV-HKU4, Uniprot ID: P0C6W3, Bat CoV HKU4; HCoV-OC43, Uniprot ID: P0C6X6, Human CoV OC43; HCoV-HKU1, Uniprot ID: P0C6X2, Human CoV HKU1; MCoV-JHM, Uniprot ID: P0C6Y0; Murine CoV (strain JHM); BCoV-HKU9, Uniprot ID: P0C6W5, Bat CoV HKU9; HCoV-NL63, Uniprot ID:P0C6X5, Human CoV NL63; HCoV-229E, Uniprot ID: P0C6X1, Human CoV 229E; BCoV-2006, Uniprot ID: S5YAF0, Bat CoV CDPHE15/USA/2006; PCoV-CV777, Uniprot ID: P0C6Y4, Porcine epidemic diarrhea CoV (strain CV777); FCoV, Uniprot ID: Q98VG9, Feline CoV; TCoV, Uniprot ID: B3FHU3, Turkey enteric CoV; AIBV-M41, Uniprot ID: P0C6Y3, Avian infectious bronchitis virus.

**Fig. S3**

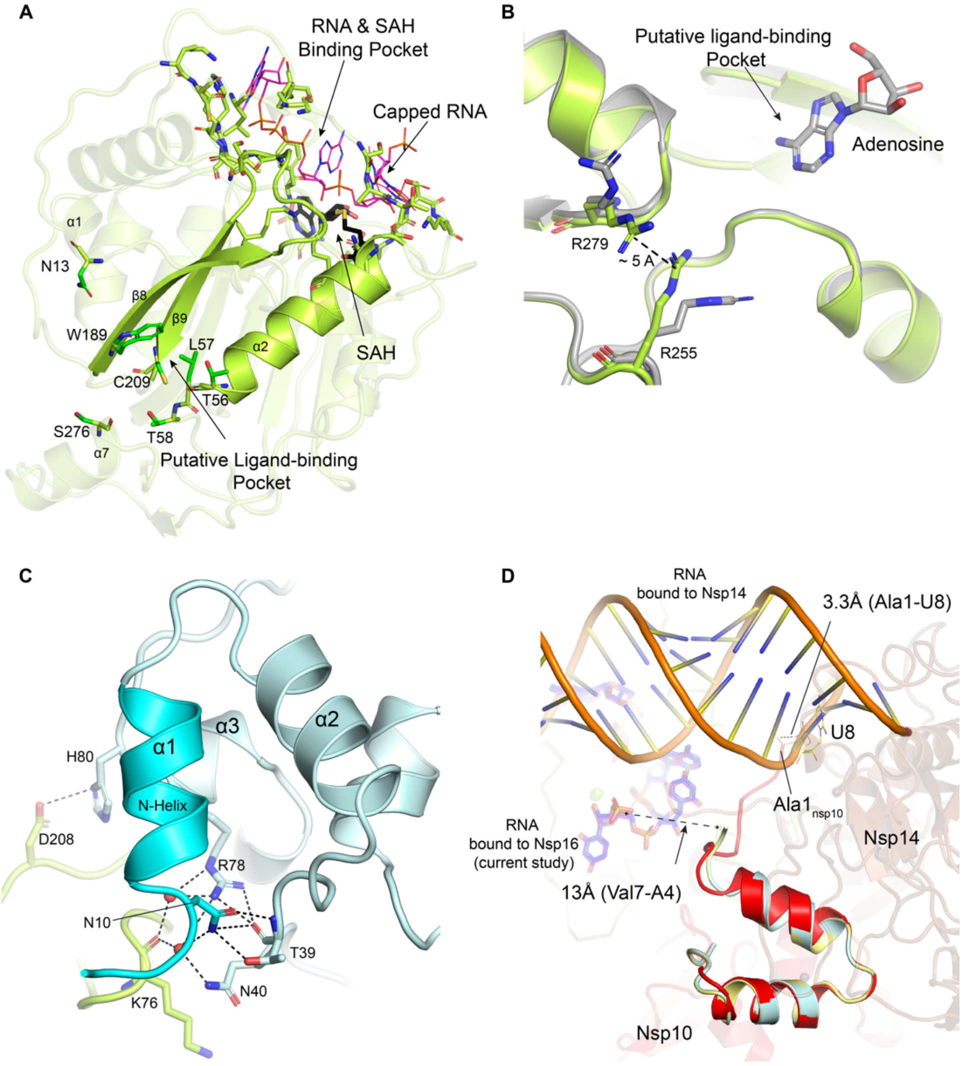


**Supplementary Fig. 3. A.** Secondary structural elements form a nonspecific nucleotide-binding site on the back side of nsp16 and the catalytic surface on the front side of nsp16. **B.** Flipped Arg side chains near the putative ligand-binding pocket (Grey, Cap-0 analog bound structure, 6WKS, and green, Cap-1 $N_{10}$-RNA bound, current study). **C.** Nsp10 N-helix, a fold-back conformation is stabilized by a network of h-bonds from the side chains of Asn10, Thr39, Asn40, and Arg78 from nsp10 and a water-mediated h-bond with Lys76 of nsp16. **D.** An overlay of the RNA-bound nsp10 in the current nsp16/nsp10 structure (nsp10: cyan and yellow) with the RNA-bound nsp14/nsp10 structure (7N0B, nsp10: red).
